# A look beyond the QR code of SNARE proteins

**DOI:** 10.1101/2024.04.24.590896

**Authors:** Deepak Yadav, Aysima Hacisuleyman, Mykola Dergai, Dany Khalifeh, Luciano A. Abriata, Matteo Dal Peraro, Dirk Fasshauer

## Abstract

Soluble N-ethylmaleimide-sensitive factor Attachment protein REceptor (SNARE) proteins catalyze the fusion process of vesicles with target membranes in eukaryotic cells. To do this, they assemble in a zipper-like fashion into stable complexes between the membranes. Structural studies have shown that the complexes consist of four different helices, which we subdivide into Qa-, Qb-, Qc-, and R-helix on the basis of their sequence signatures. Using a combination of biochemistry, modeling and molecular dynamics, we investigated how the four different types are arranged in a complex. We found that there is a matching pattern in the core of the complex that dictates the position of the four fundamental SNARE types in the bundle, resulting in a QabcR complex. In the cell, several different cognate QabcR-SNARE complexes catalyze the different transport steps between the compartments of the endomembrane system. Each of these cognate QabcR complexes is compiled from a repertoire of about 20 SNARE subtypes. Our studies show that exchange within the four types is largely tolerated structurally, although some non-cognate exchanges lead to structural imbalances. This suggests that SNARE complexes have evolved for a catalytic mechanism, a mechanism that leaves little scope for selectivity beyond the QabcR rule.

## Introduction

Eukaryotic cells are subdivided into numerous membrane-bound organelles, each enjoying a distinct microenvironment that enables them to carry out a particular function separate from other organelles. These organelles, such as the nuclear envelope, the endoplasmic reticulum (ER), the Golgi apparatus, lysosomes, vacuoles, endosomes, and the cell membrane, form a vast network known as the endomembrane system. Each organelle of the endomembrane system must maintain its identity but, at the same time, needs to exchange material with other organelles. This exchange is achieved by vesicular transport. The molecular machineries involved in this vital process are highly conserved among all eukaryotes. Cargo-laden vesicles bud from a donor compartment with the help of coat proteins, are shuttled along cytoskeletal tracks by motor proteins, and are attached to the appropriate acceptor compartment by tethering proteins. The central players in the final process of vesicle fusion, during which the cargo is released into the target organelle, are the <u>S</u>oluble <u>N</u>-ethylmaleimide-sensitive factor <u>A</u>ttachment protein <u>RE</u>ceptor (SNARE) proteins (reviewed in (Bock 2001,Jahn 2006,Jahn 2024,Rothman 1994,Südhof 2009)). These tail-anchored membrane proteins operate via a fundamental mechanism: their sequential assembly into tight membrane-bridging complexes pulls the two membranes together. Typically, four different SNARE helices assemble into a parallel four-helix bundle (Fasshauer et al. 1998; Sutton et al. 1998). Pioneering studies using a simple liposome preparation by the group of Jim Rothman showed that SNARE complex formation provides the energy that drives membrane fusion (Weber 1998). Although SNARE proteins are the driving force for fusion, their activity is orchestrated by various other factors including conserved protein families such as Sec1/Munc18 (SM), Rab, and Complexes Associated with Tethering Containing Helical Rods (CATCHR) proteins (Bonifacino & Glick 2004; Cai et al. 2007; Jahn & Fasshauer 2012; Rizo 2022; Stanton & Hughson 2023).

Right after the discovery of SNARE proteins in the early 1990s (Söllner et al. 1993), it was suggested that only certain matching combinations of SNAREs would form a proper SNARE complex that leads to membrane fusion. A matching combination, so the idea, would consist of one SNARE protein (called v-SNARE) residing exclusively on the transport vesicle and two or more SNARE proteins (called t-SNAREs) on the target membrane (Rothman 1994). Indeed, when various yeast SNARE proteins were tested in a liposome fusion assay, fusion occurred almost exclusively when cognate SNARE combinations, which were known to work in different trafficking steps, were used (McNew et al. 2000; Parlati et al. 2000, 2002; Paumet et al. 2004; Volchuk et al. 2004; Weber et al. 1998). However, promiscuous SNARE interactions were detected *in vitro* (Fasshauer et al. 1999; Furukawa & Mima 2014; Izawa et al. 2012; Wiederhold et al. 2010; Yang et al. 1999) and *in vivo* (Antonin et al. 2000; Bethani et al. 2007; Hohenstein & Roche 2001; Kreykenbohm et al. 2002; Kweon et al. 2003; Tsui & Banfield 2000; Tsui et al. 2001; Xu et al. 2014). It is therefore debated intensively whether correct assembly of SNARE proteins contributes to the specificity of membrane trafficking (for reviews see (Jahn & Scheller 2006; Koike & Jahn 2022; Linial 1997; Pelham 2001; Wendler & Tooze 2001)). It is unclear to which extent the interacting helices of SNARE proteins, the so-called SNARE motifs, are able to discriminate between cognate and non-cognate SNARE partners during complex formation.

The SNARE-motif is a conserved α-helical stretch about 60 amino acids long. In the core of the tightly packed SNARE bundle, 16 mostly hydrophobic layers are formed by interacting side chains from each of the four α-helices (Fasshauer et al. 1998; Sutton et al. 1998; Weimbs et al. 1997, 1998). The layer in the center of the complex, the so-called “0-layer”, stands out as it consists of three glutamine (Q) residues and one arginine (R) residue. The asymmetric “0-layer” is enclosed by more symmetric leucine-zipper-like layers. Along the extended SNARE bundle, several asymmetric layers exist (Fig. 1A & Fig. S1). This complementary architecture, as it probably increases the stability of the complex, might be beneficial for the catalytic activity of SNARE proteins, which is to exert the force required for membrane fusion.

**Fig. 1.**
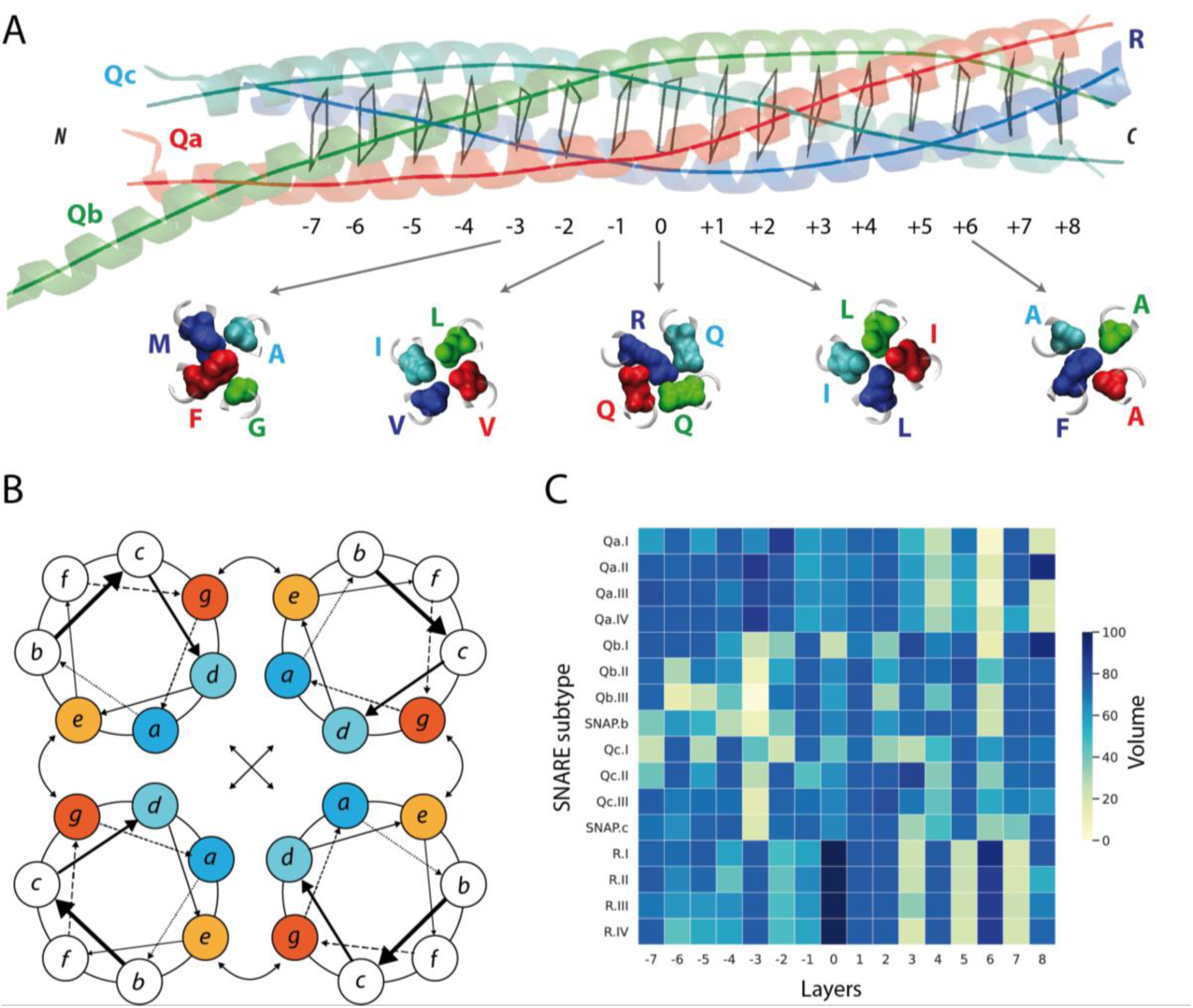
Matching pattern of large and small side chains in the core of the SNARE bundle. A) SNARE proteins form a parallel coiled coil bundle composed of four different helices. The mostly hydrophobic core of the heterotetramer contains symmetrical and asymmetrical layers. The layer in the center of the complex, the so-called “0-layer”. The complementary nature of this asymmetric layer is reflected in the strong hydrogen bonds between the guanidino group of the arginine side chain and the glutamine residues. The hydrophobic amino acids of the “0-layer” are almost unchanged throughout the SNARE protein family. The surrounding -1 and +1 layers are in ’a’ position and contain mostly leucine and valine residues. The archetypical structure of the neuronal SNARE complex (PDB: 1SFC) is shown as ribbon diagram on the top (blue, red, and green for synaptobrevin 2, syntaxin 1a, and SNAP-25a, respectively). The 16 layers (-7 until 8) in the core of the bundle are indicated by virtual bonds between the corresponding Cα positions. The structures of selected layers are shown in detail below. Additional layer structures are depicted in Fig. S1. B) Helical wheel diagram of a parallel four-helix bundle. The residues in positions ’a’ and ’d’ are at the coiled-coil interface and form a tight-fitting core. They are surrounded by the residues in positions ’e’ and ’g’ that often interact electrostatically (indicated by arrows). Residues in positions ’b’, ’c’ and ’f’ are usually exposed to the surrounding water forming the hydrophilic coat. Drawing after (Apostolovic et al. 2010). C) Plot of the vdW volume of the residues in positions ’a’ and ’d’ in the different layers of the complex for the different subtypes of SNARE proteins. For each SNARE type, the sequence analysis was restricted to the regions that form the extended coiled-coil bundle. In asymmetric layers, small and large side chains interact with each other. The analysis was carried out using alignments of the different SNARE subtypes across all eukaryotes (Kloepper et al. 2007). These alignments are shown as a weblogo presentation in Fig. S2. The VdW volume of all positions is given in Fig. S3.

Among SNARE proteins, four basic types, namely Qa-, Qb-, Qc-, and R-SNAREs, can be distinguished by their sequence profiles (Kienle et al. 2009; Kloepper et al. 2007). They correspond to the four different helices of the complex. In eukaryotes, more than 20 SNARE subtypes with unique sequence profiles exist. It is thought that they are localized to different organelles where they assemble into complexes that work in distinct vesicle trafficking steps. Their phyletic distribution among eukaryotes suggests that the SNARE subtypes represent the original repertoire of the Last Eukaryotic Common Ancestor (LECA), which was a fairly sophisticated cell with all compartments of the endomembrane system (Dacks et al. 2016; Gabaldón 2021; Koonin 2010). The LECA is thought to have lived about two billion years ago and the major eukaryotic lineages probably diverged rapidly afterwards. Remarkably, the basic set of SNARE subtypes has been maintained in almost all extant eukaryotes (Kloepper et al. 2007), but it has not been explored yet whether the two billion years of independent evolution has left marks on the interaction surfaces of SNARE complexes working in different trafficking steps.

Here, we examined the sequences of SNARE proteins to determine whether they exhibit patterns that dictate their SNARE interactions. To analyze this structurally, we exploited the physicochemical properties of the amino acids of the SNARE motif. We have modeled a wide variety of SNARE complex compositions and studied them with molecular dynamics (MD) simulations. Our analyses show that the four basic types of SNARE proteins possess an ingrained sequence pattern that assigns them their position in the “QabcR” bundle. However, the sequences do not exhibit a clear pattern that would hinder exchange within the equivalent SNARE type working in other trafficking steps. Indeed, our biochemical and MD experiments show that several exchanges within the equivalent SNARE type are possible in between trafficking steps. However, not all exchanges are possible, suggesting that some SNARE combinations are preferred.

## Results

In earlier phylogenetic studies, we had identified approximately 20 basic subtypes within the eukaryotic SNARE protein family and had developed specific as well as sensitive hidden Markov models (HMM) for each subtype (Kienle et al. 2009; Kloepper et al. 2007). In order to take a fresh look at the specificity of SNARE proteins, we made use of our large collection of about 18,000 unique SNARE protein sequences from a broad range of eukaryotes. We classified the sequences into the four basic types, namely Qa-, Qb-, Qc-, and R-SNAREs. By means of phylogenetic studies, these basic types could be further divided into subtypes with specific sequence profiles. These subtypes correspond to the SNARE proteins assigned before to different cellular transport steps based on cell biological analyses, i.e. SNARE proteins involved in the following trafficking steps: ER (I), Golgi (II), endosomes (III), and secretion (IV).

### From sequence to amino acid properties

In a coiled-coil, each helix is stabilized by a regular pattern of hydrogen bonds between the peptide N-H and C=O groups, while the helices are held together by hydrophobic interactions in the core (Woolfson 2005,Mason 2004,Grigoryan 2008,Lupas 2017,Truebestein 2016,Woolfson 2023,Schreiber 2011,Zhang 2009,Apostolovic 2010). In the regular heptad repeats, labeled ’abcdefg’, the ’a’ and ’d’ positions interact as leucine-zipper-like layers in the interior (Fig. 1B). The conservation pattern of each SNARE subtype showed, in agreement with earlier observations (Fasshauer et al. 1998; Weimbs et al. 1998), that the residues that form the 16 coiled-coil layers are usually highly conserved (Fig. S2). The 16 coiled-coil layer residues, except for the ionic 0-layer, are mostly hydrophobic across all different SNARE subtypes (Fig. S3a). Besides the conspicuous R- and Q-residues of the 0-layer, which possibly aligns the four helices, the hydrophobicity plot did not provide novel insights into the selectivity of SNARE proteins.

Along the extended SNARE bundle, several asymmetric layers exist in which bulky side chains are packed together with smaller ones (Fig. 1A & Fig. S1). Structures of different SNARE complexes have revealed that the overall structure and the arrangement of layers is conserved. When we plotted the van der Waals volume (vdW) of the side chains a more diverse pattern appeared (Fig. S3b). When only the vdW volume of the side chain in ’a’ and ’d’ positions was plotted, an intriguing matching pattern of large and small side chains in several layers of the bundle becomes apparent (Fig. 1C), suggesting that packing complementarity of side chains in the core of the bundle could be key for tight SNARE complex formation.

### From amino acid properties to 3D models

To test this hypothesis, we looked at the volume of side chains in different layers of SNARE complexes. The structures of available SNARE complexes are highly similar, with root mean square deviations (RMSD) between 0.9 to 2.3 Å (Diao et al. 2015; Strop et al. 2008), implying that the overall structure of SNARE complexes has been largely maintained during evolution. The four helices in the bundle are highly twisted, but an overall pitch cannot be defined, as each helix has its own unique character. Interestingly, the radius of SNARE complexes changes over the length of the bundle, giving it a spindle-like shape. The largest expansion can be seen in the center of the complex. The central 0-layer is surrounded by layers with large hydrophobic amino acids that seem to seal the hydrophilic 0-layer. Adjacent layers on both sides, i.e., towards the *N*-terminal portions and towards the membrane anchors on the *C*-terminal side, tend to occupy less volume. As outlined before, these layers are often asymmetric and combine voluminously with small side chains.

In lack of a simple geometrical description of hydrophobic core of SNARE complexes, we calculated the volumes of virtual cubicles between Cα atoms of amino acids in ’a’ and ’d’ positions to inspect the contribution of each helix to the complementarity of the hydrophobic core. We assumed that the side chains of layer positions occupy about half of the volume of each of two neighboring cubicles (Fig. 2A). When we plot the average volume of two neighboring cubicles, the spindle-like shape of the volume distribution in SNARE complex structures becomes visible (Fig. 2B). For each complex, we then calculated how much space is occupied by the side chains of layer residues. The volume occupied by the side chains also showed a spindle-like distribution with additional minima in layers -3, 3 and 6 (Fig. 2B). For a first overview of all SNARE proteins, we combined classified sequences within different subtypes into SNARE complexes. Each of these hypothetical SNARE complex combinations was composed of four basic types, Qa-, Qb-, Qc-, and R-SNAREs. When we calculated the volumes of ’a’ and ’d’ positions in these presumed SNARE complexes across eukaryotes, the distribution was comparable to the ones obtained from X-ray structures (Fig. 2B), suggesting that SNARE complexes indeed have a common design.

**Fig. 2.**
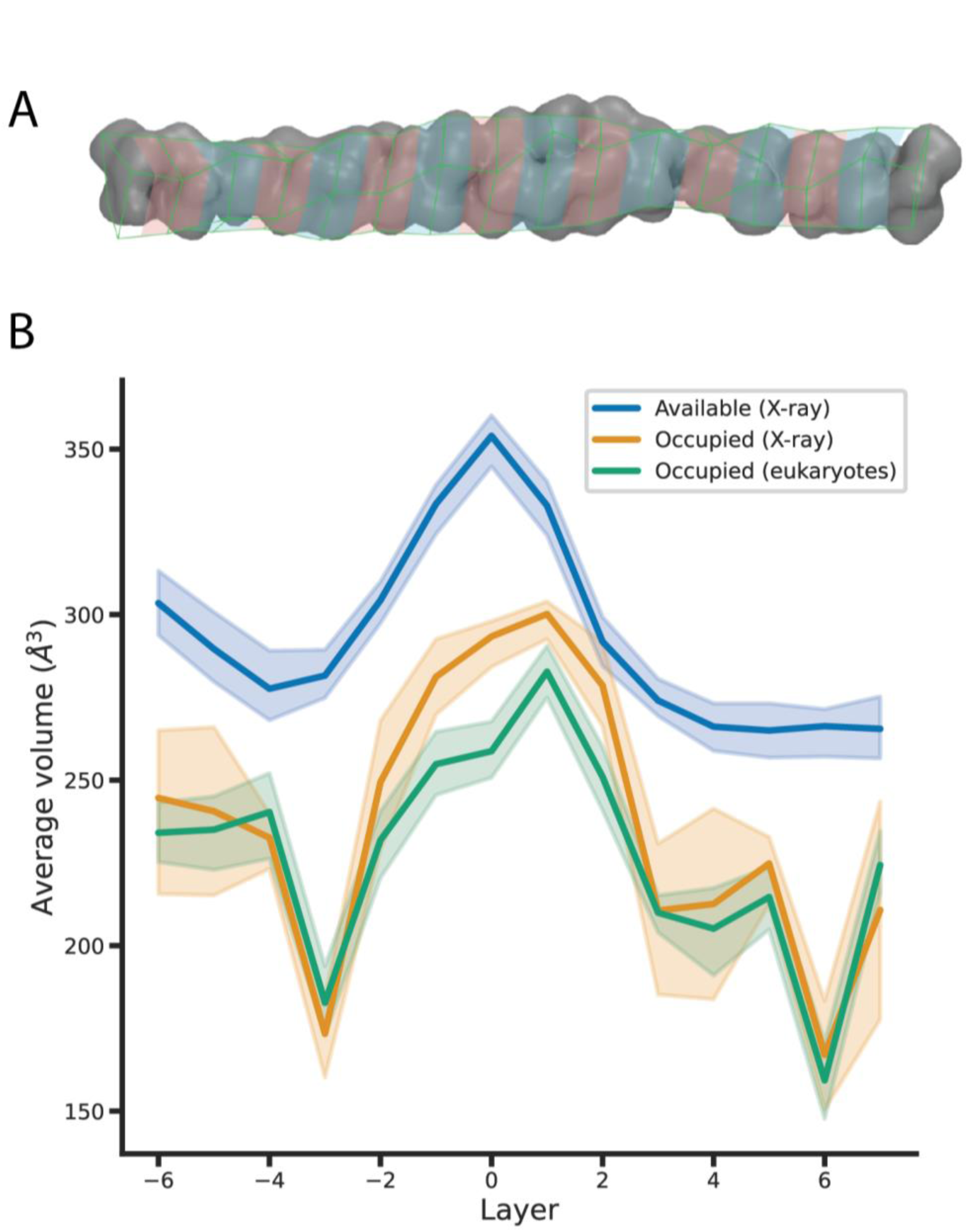
Conserved spindle-like shape of SNARE complexes. The volume available and the volume filled by amino acids in the core of the SNARE bundle changes in a spindle shape-like manner. The available volume of the hydrophobic core was inferred from virtual cubicles between Cα atoms of amino acids in ’a’ and ’d’ registers of the SNARE motifs (A). The largest volume is available to the 0-layer with its four voluminous side chains. The available volume and the occupied volume were calculated using existing crystal structures of SNARE complexes. For the calculation, cubicles were introduced between the Cα atoms of amino acids in ’a’ and ’d’ positions as shown (B). In order to calculate how much volume other SNARE combinations would occupy, the volumes of amino acids in ’a’ and ’d’ positions from Qa-, Qb-, Qc- and R-SNAREs were added, revealing that the shape of the SNARE core is largely conserved across eukaryotes.

To gain more insights into the question of whether SNARE complexes must be assembled according to a ’QabcR rule’, i.e. that they must be composed of the four basic SNARE types, we computed the side chain volumes of the layer residues for combinations of four SNARE proteins from baker’s yeast within the different trafficking steps, including those combinations that do not consist of four different helices. For example, baker’s yeast has four SNARE proteins working in trafficking towards the ER, Ufe1 (Qa.I), Sec20 (Qb.I), Use1 (Qc.I), and Sec22 (R.I). If we consider all potential arrangements of these four proteins involved in ER trafficking step, including the possibility of having multiple proteins of the same SNARE type in the complex, there are 35 different combinations. There are even more combinations of yeast SNARE proteins from other trafficking steps. Most of them deviating from the ’QabcR rule’, i.e. including all hypothetical combinations that contain one or more of the same SNARE type (Fig. S4).

Combinations of four yeast SNARE proteins following the ’QabcR rule’ fit well into the hydrophobic core of the spindle-like complexes, whereas many other combinations did not (Fig. S4). Although the complexes which follow the ‘QabcR rule’, QabcR-complexes, always performed well, this approach was unable to discriminate strictly between QabcR-complexes and several other combinations (Fig. S4). However, when we subdivided the SNARE proteins according to the different vesicle transport steps and restricted the analysis to parallel complexes, the predictive power improved somewhat: QabcR complexes performed best (Fig. 3), corroborating the notion that a combination of four different SNARE types produce a compact interaction through complementary side chain packing.

**Fig. 3.**
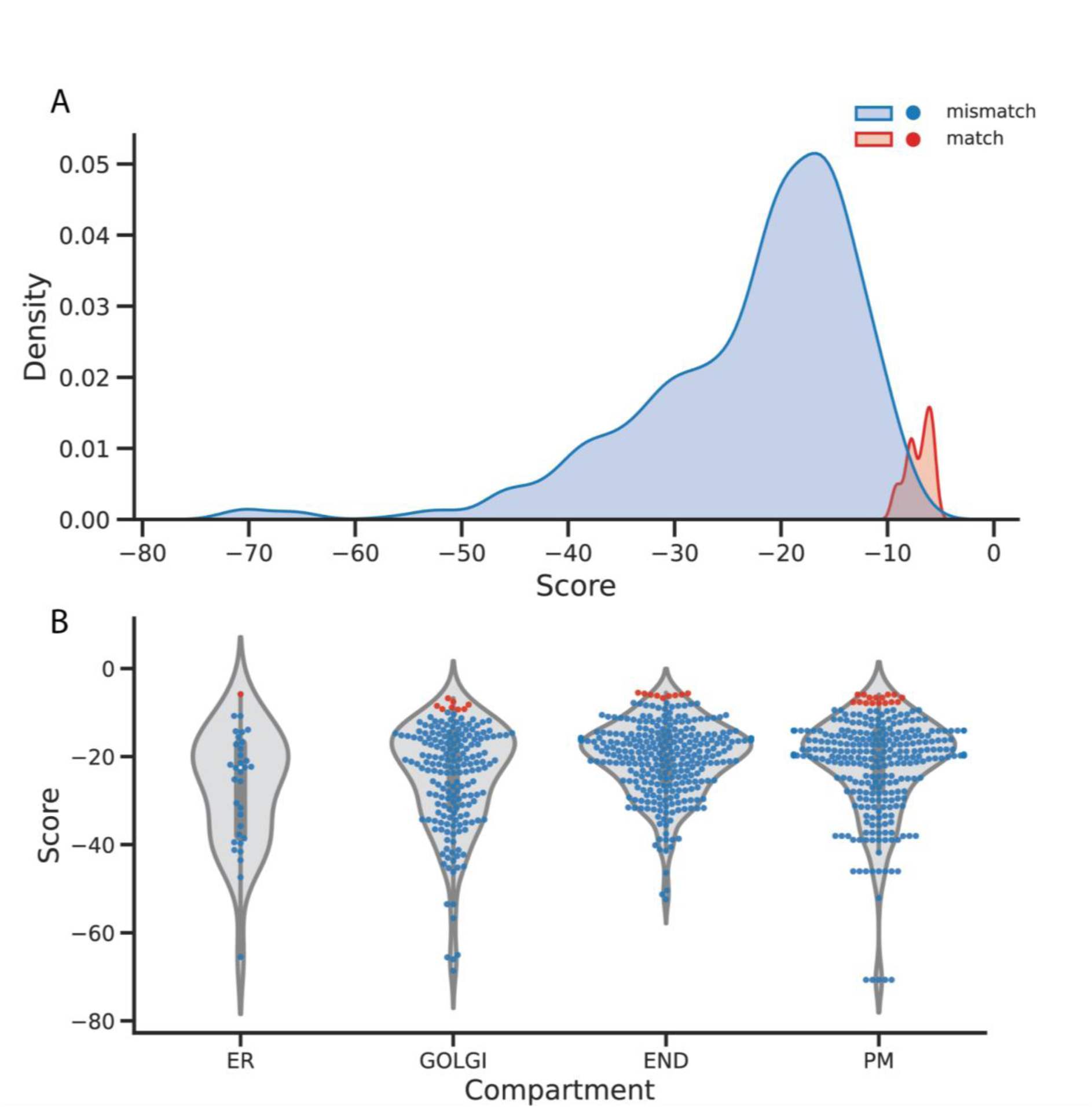
Combinations of four different snare helices following the ’QabcR rule’ fit best into a spindle-shaped complex. SNARE proteins from baker’s yeast were assembled in different combinations, but in the respective transport types, ER (I), Golgi (II), endosomes (III), and secretion (IV). Note that the subtype III included SNAREs that are thought to work in trans Golgi network (TGN), endosomal and lysosomal trafficking steps. For each combination, the volume of the core was calculated from the VdW of the amino acids in ’a’ and ’d’ positions. While the volume of established QabcR combinations shows the familiar spindle-shaped pattern, other combinations deviate significantly from this pattern (A). However, a clear separation of all QabcR combinations (red) from other combinations (blue) was not achieved. A better separation of QabcR combinations from other combinations was achieved if only the available SNARE combinations of a subtype were considered (B). When ordered by transport steps, the QabcR combinations scored best.

### Different physico-chemical properties of the four SNARE helices

The above analyses confirm that the physico-chemical properties of the side chains, hydrophobicity and vdW volume, of SNARE proteins play a significant role for the correct assembly of the four helices. We therefore included other physicochemical descriptors of side chains (Abriata et al. 2015; FAUCHÈRE et al. 1988), for our analysis, viz., flexibility, bulkiness, steric factor and maximal number of hydrogen bonds a side chain can establish. A principal component analysis (PCA) was performed on these six physicochemical descriptors and the first two principal components were used to visualize the proteins. With this approach, we were able to discriminate between the four major SNARE types. In particular, most R-SNAREs were well separated from all other SNARE types, whereas the boundary between Q-SNAREs was somewhat less distinct (Fig. 4). This is consistent with phylogenetic studies that have shown that Q-SNAREs are generally less distinct compared to R-SNAREs (Kloepper et al. 2007; Neveu et al. 2020). Overall, this analysis indicates that each of the four basic SNARE types is endowed with a characteristic physicochemical profile. We were wondering whether these different physicochemical profiles could guide the four SNARE types into the correct helix position in the four-helix bundle? To better understand this, we have studied structures of SNARE complexes in more detail.

**Fig. 4.**
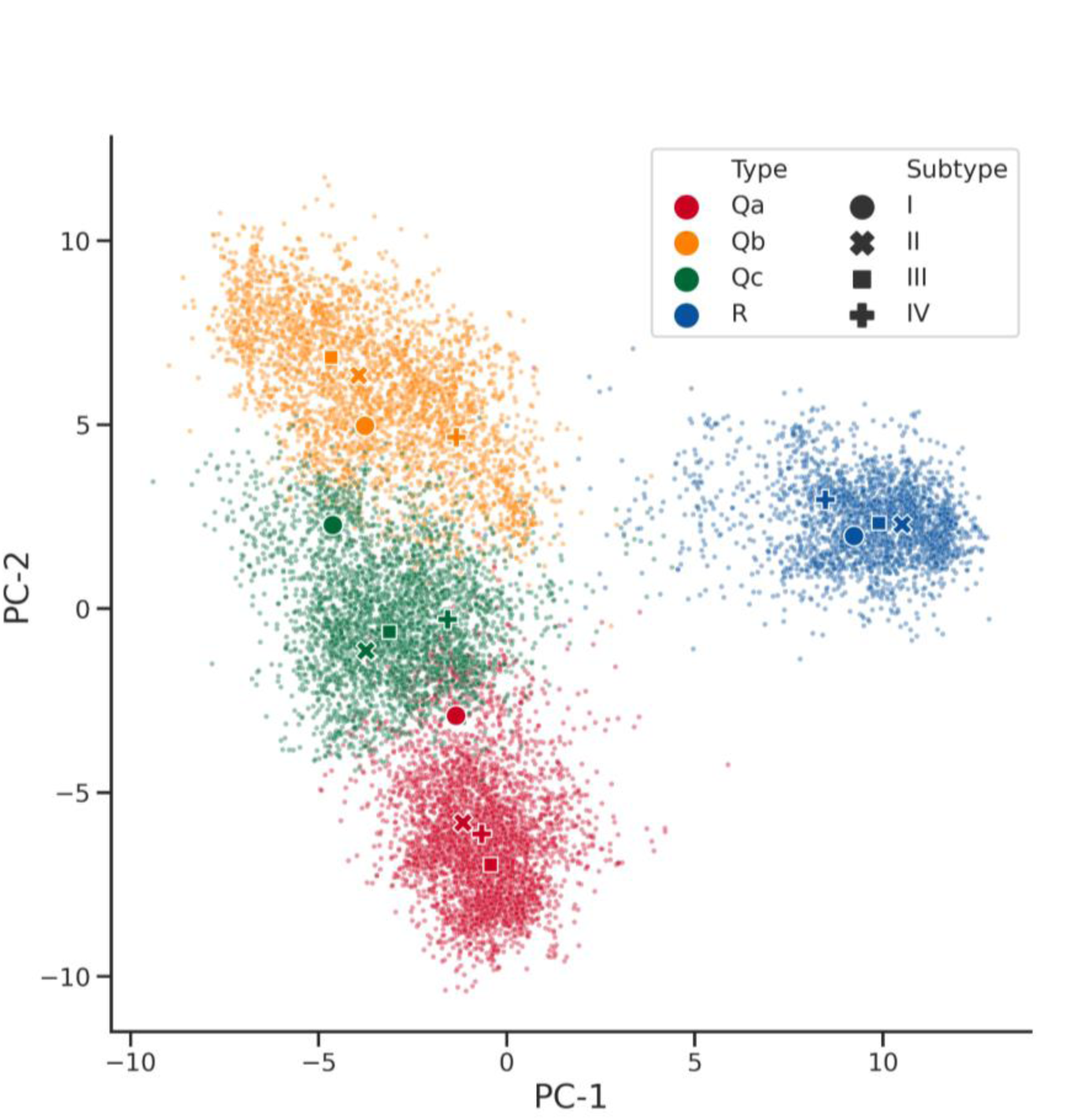
The four SNARE types can be distinguished on the basis of their amino acid composition. Visualization of principal component analysis (PCA) of SNARE protein amino acid compositions based on the physico-chemical properties of their side chains. The PCA was performed on six physicochemical descriptors, hydrophobicity, vdW volume, flexibility, bulkiness, steric factor and maximal number of hydrogen bonds a side chain can establish. The first two principal components were used to visualize the proteins. The plot shows that R-SNAREs (blue) in particular have their own profile and are clearly separated from all other types. Qa- (red) and Qb-SNAREs (dark yellow) are also separated, but partly overlap a little with Qc-SNAREs (green), which lie between the two types. The overlap is mainly due to Q-SNAREs involved in the ER transport step (i.e., type I). These SNARE proteins of type I are in fact more divergent.

In the analyses above, we focused on the side chains of the amino acids that form the hydrophobic core of the four-helix bundle. Structures of SNARE complexes show that the ’a’ and ’d’ positions have the highest number of contacts, followed by ’e’ and ’g’ positions, whereas ’b’ and ’c’ have only few contacts and ’f’ positions do hardly interact with other residues (Fig. S5). When we plotted the physicochemical descriptors for each of the different positions of the heptad repeat independently, ’a’ and ’d’ positions, which form the core layers, contributed most. ’e’ and ’g’ positions also contributed significantly, whereas ’b’, ’c’ and ’f’ residues, which point to the outside of the complex, did not contribute much to the divergent profiles of the basic SNARE types.

### SNARE complex models

The results obtained so far suggest that SNARE proteins preferentially assemble into bundles composed of four different helices. However, this analysis does not allow any conclusion regarding the structural arrangement of the four different helices in the bundle. To gain deeper insights into this, we used the crystal structures of SNARE complexes, 1sfc, 1gl2, 2nps, and 3b5n, as a basis for structural permutations. For each complex, 70 different permutations were possible, ranging from four different helices to four helices of the same kind in one complex.

For each permutation, we used the Qa-, Qb-, Qc- and R-helices from the respective crystal structures and the sequences were threaded onto the structure using Modeller. Each model was then optimized using the CoupledMoves protocol of Rosetta (Ollikainen et al. 2015). Side-chains and backbones were allowed to move. Top three decoys were selected from CoupledMoves, and the performance of every decoy of each model was evaluated using three parameters: i) overall energy scores, ii) presence of intramolecular void volumes, and iii) atom-to-atom contacts.

When these parameters were plotted for all permutations for a given SNARE set, the combination QabcR scored best among all permutations (Fig. 5 & Fig. S6). This suggests that other combinations fit less well into the architecture of the SNARE bundle, probably caused by mismatches of residues of the hydrophobic core. This supports our initial idea that the inward pointing side chains of QabcR helices interlock into a tight-fitting hydrophobic core. Interestingly, the circular arrangement Qa→Qb→Qc→R→ scored best among all six possible permutations of four different helices in the same complex, suggesting that this arrangement is structurally preferred. In other words, the position of each helix in the four-helix bundle appears to be fixed.

**Fig. 5.**
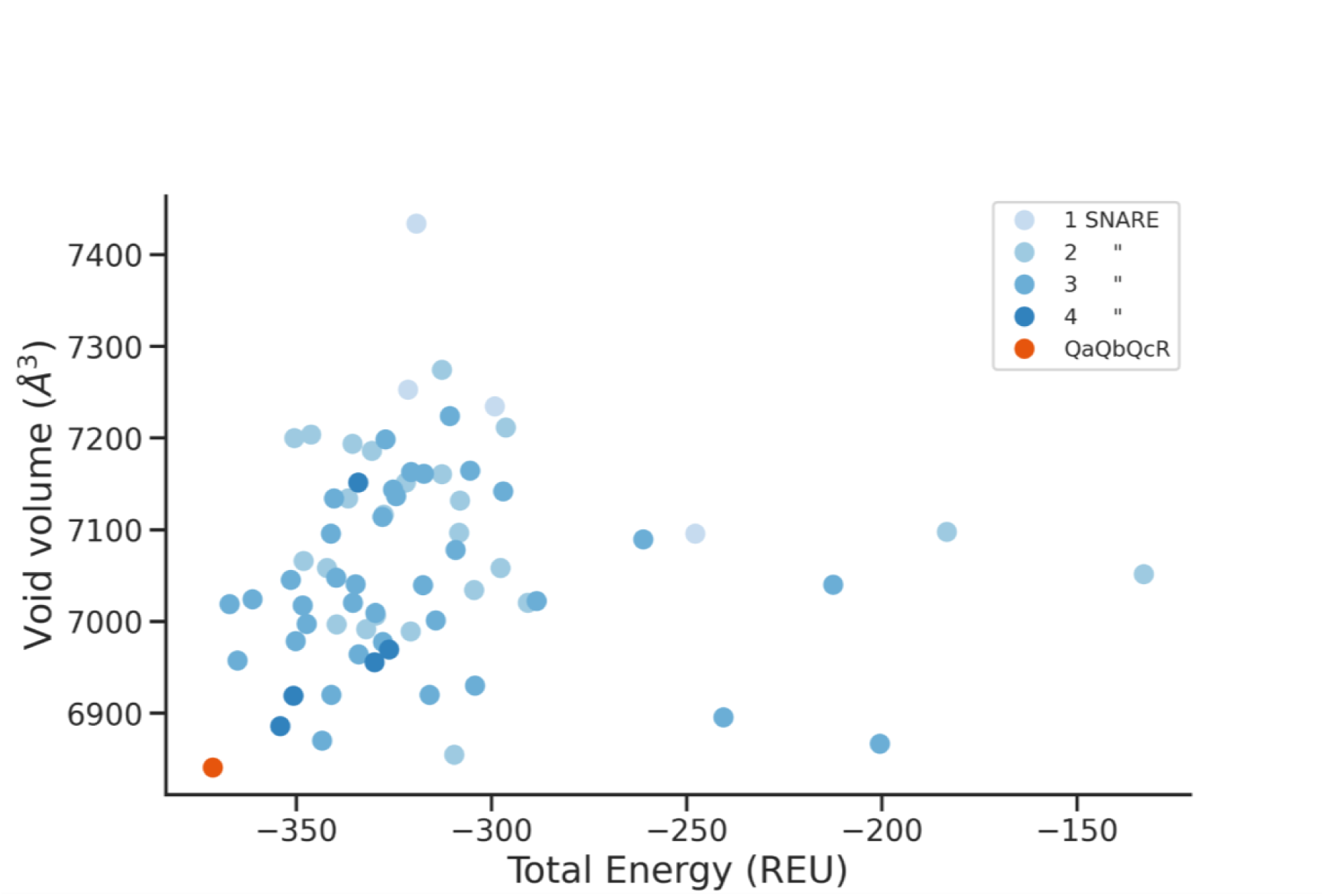
Complexes with a QaQbQcR arrangement of the four helices are energetically favored. 70 different structural permutations of the neuronal SNARE complex were modeled and their performance was evaluated using three parameters: i) overall energy scores, ii) presence of intramolecular void volumes, and iii) atom-to-atom contacts. When plotting the void volume against the overall energy, the arrangement of the helices as present in the crystal structure (pdb: 1sfc) is the most favorable. A similar result is obtained when the atom-to-atom contacts were plotted against the overall energy (Fig. S6A). The corresponding results for other SNARE complexes (pdb: 1gl2, 3b5n, 2nps) are shown in Fig. S6B-D.

### Biochemical analysis of cognate and non-cognate SNARE complexes

A number of biochemical studies have shown that SNARE proteins are capable of forming non-cognate complexes as long as the QabcR rule is followed (Fasshauer et al. 1999; Jahn & Scheller 2006; Khalifeh et al. 2022; Linial 1997; Neveu et al. 2020; Pelham 2001; Tsui & Banfield 2000). However, it is still not clear whether all QabcR combinations can form stable complexes. Obviously, it would be a very large effort to perform such analysis exhaustively, and we therefore decided to use two divergent cognate SNARE complexes, the SNARE complex involved in neuronal secretion and the SNARE complex involved in trafficking towards the ER for selected biochemical tests. The neuronal SNARE complex is composed of syntaxin 1a (Syx1a, Qa.IV), SNAP-25 (Qbc.IV), which is composed of two SNARE domains of different subtypes, SN1 (Qb.IV) and SN2 (Qc.IV), and synaptobrevin 2 (Syb2, R.IV). The ER SNARE complex is composed of syntaxin 18 (Syx18, Qa.I), Sec20 (Qb.I), Use1 (Qc.I), and Sec22 (R.I), but is much less well studied biochemically. When we mixed the respective purified proteins both complexes formed (Fig. 6). We then started various exchange experiments. In the first set of mixing experiments, Qa-SNAREs in the ER and neuronal SNARE complex were exchanged with Qa-SNAREs from different intracellular membrane trafficking pathways. More precisely, Syx18 in the ER-SNARE complex was exchanged for Syntaxin 5 (Syx5, Qa.II-), Syntaxin 16 (Syx16, Qa.III-), or Syx1a (Fig. 6b), while Syx1a in the neuronal SNARE complex was exchanged with Syx18, Syx5, (Fig. 6a) or Syx16 (Fig. S7a). With one exception, non-cognate SNARE complexes were observed using non-denaturing PAGE and size-exclusion chromatography (Fig. 6): we could not detect a stable complex when Syx1a was incubated with the ER-SNAREs Sec20, Use1, and Sec22 (Fig. 6b).

**Fig. 6.**
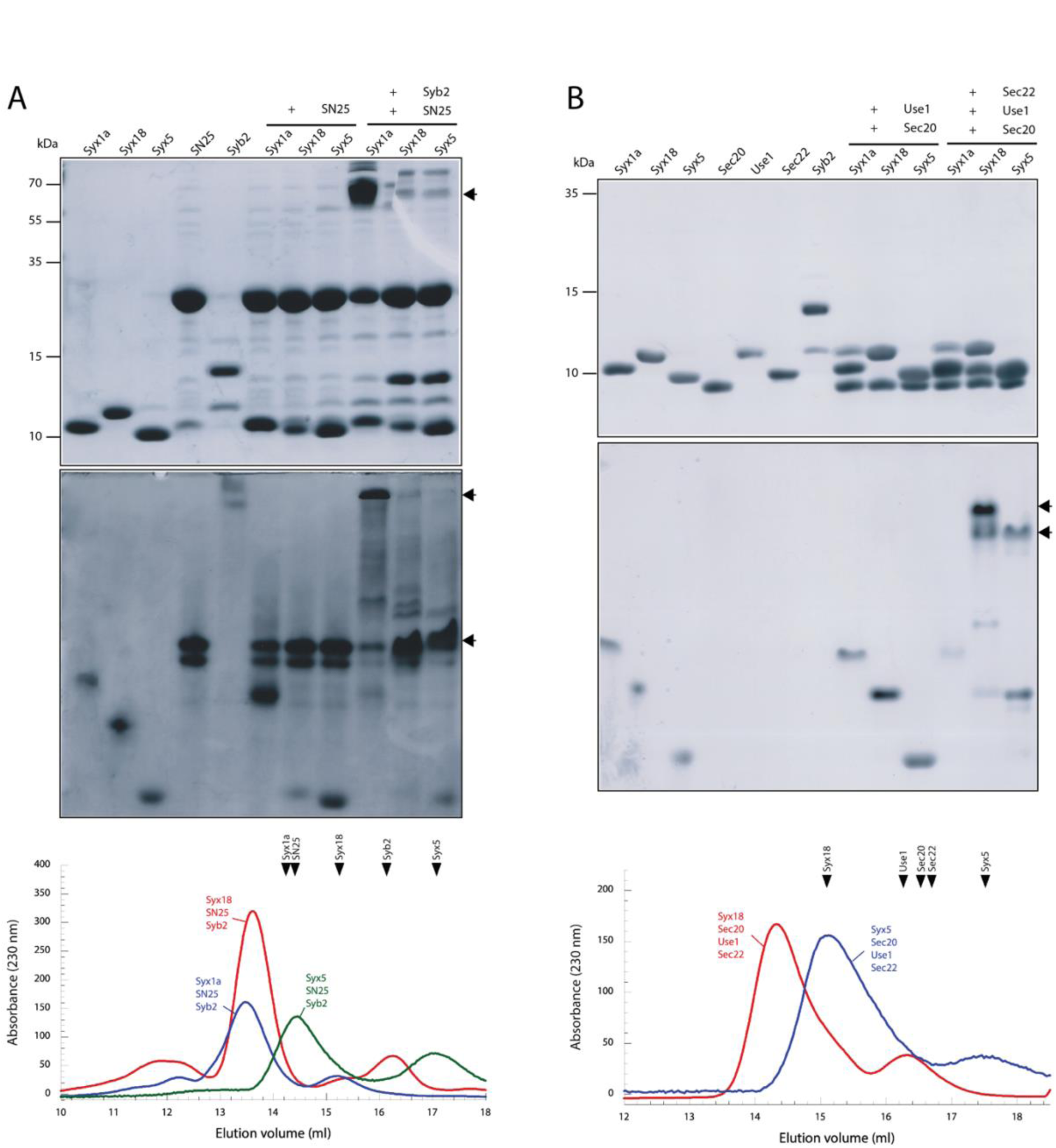
Formation of stable non-cognate SNARE complexes. The neuronal SNARE complex consists of three SNARE proteins, Syx1a (Qa), SNAP-25 (Qbc), and Syb2 (R), whereas the ER SNARE complex is composed of the four SNARE proteins Syx18 (Qa), Sec20 (Qb), Use1 (Qc) and Sec22 (R). These cognate complexes were assembled by incubating the purified monomers overnight at 4°C with equimolar ratios at ∼15 μM concentration. Both complexes were also incubated with non-cognate syntaxins (Qa) including the Golgi syntaxin Syx5. The mixtures and individual proteins of the neuronal complex **(A)** and the ER complex **(B)** were then separated by SDS-PAGE (upper panel), non-denaturing PAGE (middle panel) or size exclusion chromatography (lower panel). The band of the SDS-resistant neuronal SNARE complex in the upper panel of **(A)** is indicated by an arrow. Note that the ER complex is not SDS-resistant. Stable complexes in the non-denaturing gels, which separate native proteins and stable interactions by their charge only (middle panel), are also indicated by arrows. Proteins in gels were visualized by Coomassie Blue staining. SNAP-25 and Syb2 formed stable complexes with Syx18 and Syx5 as can be seen in the non-denaturing gel and upon size-exclusion chromatography. The exclusion volumes of the individual proteins are indicated on top by arrows. The chromatograms of the individual SNAREs are shown in Fig. S8. The ER SNAREs Sec20, Use1 and Sec22 formed a stable complex with Syx5 but not with Syx1a. Stable complexes also formed when Syx1a was exchanged with Syx16 or Syb2 with Sec22 in the neuronal complex led to stable SNARE complexes (Fig. S8).

In a second set of experiments, we exchanged Qb- and Qc-SNAREs in the ER SNARE complex. To simplify the experiments, we generated chimeric Qbc-SNARE proteins that contain a Qb- and a Qc-SNARE connected by the flexible linker region of SNAP-25. These chimeras mimic the domain structure of the Qbc-SNARE SNAP-25. We generated Qbc-SNAREs that combine cognate pairs involved in ER (type I, Sec20/Use1) and in Golgi trafficking (type II, Membrin/Bet1; Gos28/Gs15) and non-cognate combinations (Membrin/Use1; Sec20/ Bet1). Again, we found mostly stable SNARE complexes but we could not detect a complex when the Golgi Qbc-SNARE Gos28/Gs15 was mixed with the ER-SNAREs Syx18 and Sec22 (Fig. S6e). We also noted that only a weak SNARE complex band was observed when the other Golgi Qbc-SNARE Membrin/Bet1 was mixed with the ER-SNAREs. In the last set of experiments, we exchanged Syb2 in the neuronal SNARE complex with the R-SNAREs Sec22; a stable non-cognate complex was observed (Fig. S6b). Our results show that several non-cognate combinations of SNARE proteins can form stable complexes. However, there appear to be exceptions that suggest that in certain, non-cognate QabcR combinations there are structural difficulties.

### MD analysis of cognate and non-cognate SNARE complexes

To gain a more comprehensive understanding of these structural difficulties of non-cognate SNARE combinations, we ran MD simulations of protein combinations derived from our biochemical experiments. As in the biochemical experiments above, we first examined the two cognate complexes, the neuronal complex and the ER complex. The structure of the neuronal complex is solved (1sfc) (Sutton 1998) and thus could be inspected directly by MD simulation. All MD simulations were performed as three replicas over 100 ns each as described in detail in the method section. By contrast, the structure of the ER complex is not available as a high-resolution structure. With Modeller, we generated a model of the ER complex using available structures of different SNARE complexes. Then we idealized and relaxed the model by using Rosetta. Note that all further exchanges of SNARE proteins in the neuronal and the ER SNARE complexes for non-cognate combinations were generated by the same procedure. Subsequently, we performed exchanges for both cognate complexes as described above for the biochemical experiments. In the neuronal complex, we replaced Syx1a with Syx16, Syx5, or Syx18. In the ER complex, we replaced Syx18 (Qa) with Syx16, Syx5, or Syx1a. We additionally replaced Use1 (Qc) with Bet1 and in another experiment, the Qb- and Qc- helices with the Golgi SNAREs Gos28 and Gs15, respectively.

In order to measure the effects of these exchanges on the global structure of the four-helix bundle, we first used their RMSDs, which quantitatively measures the similarity of structures. For both cognate complexes, the neuronal and the ER SNARE complex, the RMSDs were obtained from the fluctuation in MD simulations (Fig. 7A, Table S1). Exchanges of the Qa-helix in the neuronal SNARE complex led to an increase in RMSDs compared to the cognate neuronal complex. In the ER complex, similar exchanges also led to comparable increases in RMSDs. The replacements of Syx18 by Syx1 or of Sec20 and Use1 by Gos28 and Gs15, respectively, in the ER complex led to the largest deviations from the cognate complex structure (Table S1).

**Fig. 7.**
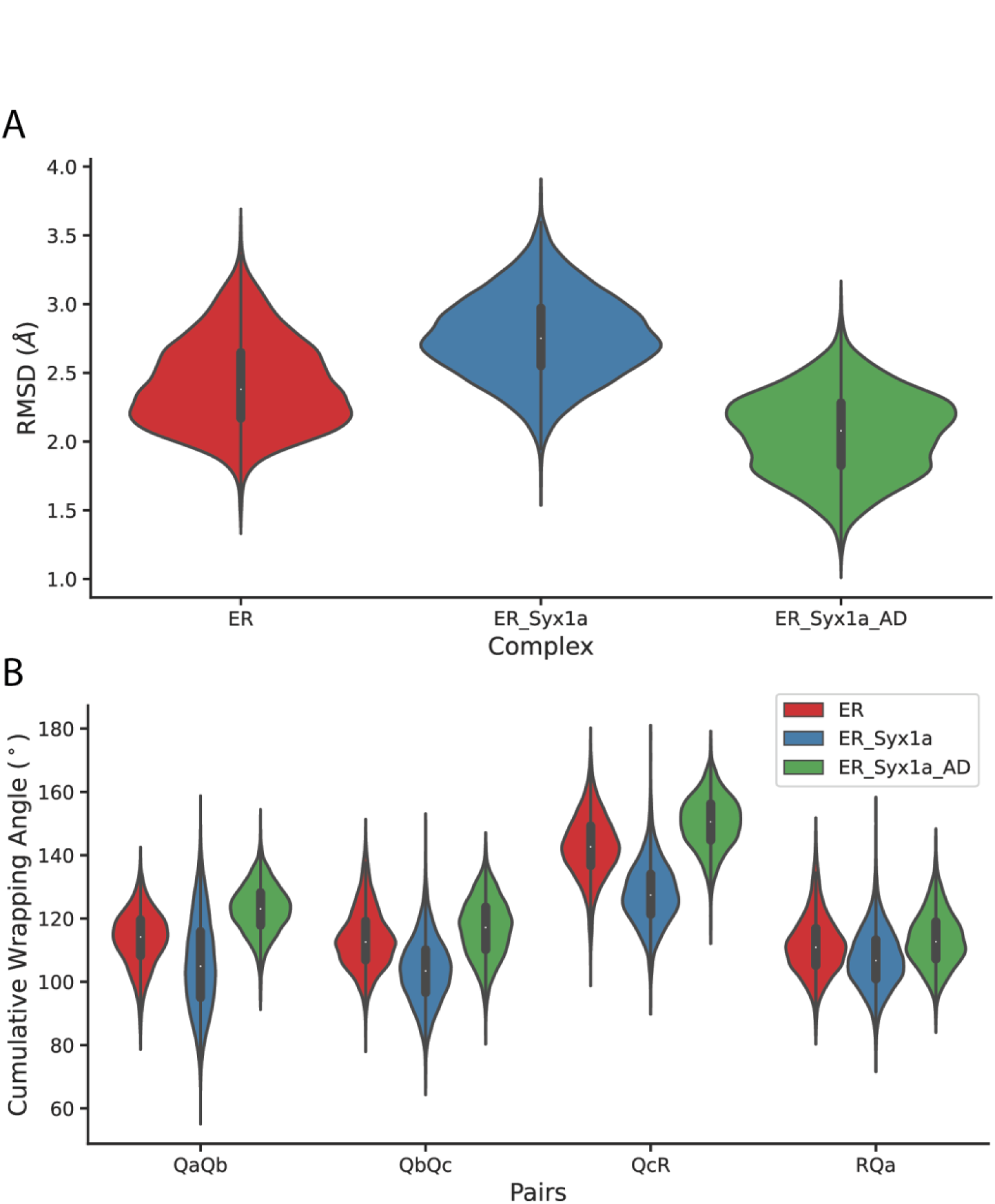
MD simulations of cognate and non-cognate ER SNARE complexes. (A) Mean RMSD values and (B) distribution of cumulative wrapping angles during MD simulations for the replacement of Syx18 with Syx1a in the ER SNARE complex. First, the simulations for the ER-SNARE complex (red) were carried out. Then for the Qa-helix Syx18 was exchanged for Syx1a (blue). Then, for the next simulation, the ’a’ and ’d’ positions of Syx1a in the ER complex were swapped back with the cognate amino acids of Syx18 (ER_Syx1a_AD, green). The results of the exchanges and MD simulations in other complexes are shown in Figs. S10 and S11.

A closer inspection of the structural fluctuations during the MD simulations revealed more drastic effects for these combinations, which were not well described by the RMSDs, however. We observed that the helices at certain regions partially detached from the association of the hydrophobic core during the simulations. These fluctuations indicate that these non-cognate complex combinations are somewhat more unstable, as suggested by the biochemical analyses above. To better describe these structural fluctuations, we then measured the movements of each of the neighboring helix pairs in the four-helix-bundles during simulations. This was achieved through calculating the six angles between the seven residues in ’d’ position throughout every helix pair, termed cumulative wrapping angle (CWA, Fig. S9). The analysis revealed that the four-helix bundle exhibited distinct distributions for CWAs of each R-Qa, Qa-Qb, Qb-Qc, and Qc-R helix pair. These distributions shifted when structure-distorting substitutions were made as shown for the exchange of Syx18 with Syx1a in the ER complex (Fig. 7B).

In addition, we measured the distances between different ’a’ and ’d’ positions in the respective complex (Fig. S11). This analysis showed even better when parts of helices were detached from the bundle during simulations.

To counteract the drastic effects of the exchanges in the ER complex, we subsequently generated models in which the ’a’ and ’d’ positions of the exchanged helices were substituted with those of the cognate helices. When we introduced Syx1a, which was provided with all side chains of Syx18 in ’a’- and ’d’-position (Syx1a_AD_), into the ER complex, the complex was more stable compared to the ER complex containing the non-cognate Syx1a helix. In contrast, the corresponding back-swap in the ER complex with Gos28/Gs15 (Gos28_Gs15_AD_) did not lead to a significantly improved fit of the hydrophobic core of the bundle. A closer inspection of the complex structure showed that this was caused by an incompatibility of the new ’a’ and ’d’ positions with the side chains of the ’e’ and ’g’ positions of the Gos28 and Gs15, suggesting that the side chains in ’e’ and ’g’ register contribute to the accuracy of fit.

In conclusion, our *in silico* approach corroborated our biochemical observations that several non-cognate SNARE combinations can form SNARE complexes. Some non-cognate combinations can lead to structural weaknesses in the four-helix bundle, however. It is worth mentioning that the non-cognate combinations that did not form a complex in the above biochemical experiments showed clear structural problems in the simulations, including the detachment of entire helical segments from the assembly.

## Discussion

The exchange of material between the compartments of the eukaryotic cell takes place via transport vesicles. It is essential for the survival of a cell that the vesicles are transported to the correct compartments. This is ensured by already providing the vesicles with the correct address, i.e., the proteins responsible for docking and fusion with the target compartment, during packaging. SNARE proteins catalyze the membrane fusion reaction after the vesicle has reached the target compartment. There, depending on the transport step, three to four different SNARE proteins form a stable heterologous four-helix bundle between the vesicle and the target membrane in a zipper-like manner, bringing together the membranes. What contribution SNARE proteins have to the specificity of this step is unclear. Based on their cellular distribution, SNARE proteins were originally divided into v- and t-SNAREs (Rothman 1994). However, this topological classification is ambiguous and with increasing structural knowledge, they have come to be referred to as Q- and R-SNAREs (Fasshauer et al. 1998). In the present work, we explored the extent to which the sequences of SNARE proteins provide insights into their interaction patterns. Can we better understand their QR code?

Our previous phylogenetic analyses had shown that all SNARE proteins can be classified into four basic types: Qa, Qb, Qc, and R (Kienle et al. 2009; Kloepper et al. 2007). These types correspond to the helices in the four-helix bundle. However, whether there is a rule for the arrangement of the four helices in the bundle has not been investigated further until now. To do so, we have now considered their physicochemical properties in our analyses. Our analyses show that the four SNARE types each have a specific physicochemical profile that enables them to form a stable four-helix bundle in a given structural arrangement. For this geometrical matching pattern, the size and chemical character of the side chains are important. Not only the central 0-layer with its hydrophilic amino acids but also some other layers of the SNARE bundle are asymmetrically built (Fig. 1a). There, space-consuming and compact side chains interact with each other, so that the bundle has a twisted spindle-like structure. We showed here that this QabcR rule, which dictates the order of the four helices in the bundle, is conserved in all SNARE complexes. It is likely that this structural design plays a role in the function of the complex as the tight interlocking could stabilize the complex during zippering and provide the necessary energy for catalyzing membrane fusion.

The four basic SNARE types can each be further divided into different subtypes based on specific sequence profiles. About 20 conserved types of SNARE proteins were present in the LECA repertoire (Kloepper et al. 2007). In many eukaryotic lineages, for example in animals and plants, the set of SNARE proteins has even been expanded significantly (Kienle et al. 2009; Kloepper et al. 2008; Sanderfoot 2007). Cells therefore have a repertoire of different SNARE subtypes available for the various transport steps between the intracellular compartments. It is thought that SNARE proteins typically form specific QabcR complexes, known as cognate complexes, during membrane fusion events within cells. Their specific pairing, so the idea, ensures that membrane fusion occurs only between appropriate membranes, preventing non-specific fusion events (Rothman 1994). Regulatory proteins contribute to the specificity of SNARE-mediated membrane fusion (Jahn et al. 2024; Koike & Jahn 2022; Linial 1997; Pelham 2001; Wendler & Tooze 2001). Despite many studies, the question of how much selectivity – beyond the basic QabcR rule – is engraved in the sequence profile of the various SNARE subtypes has remained unanswered. A clear answer is made difficult by the fact that SNARE proteins can interact promiscuously in the cell (Antonin et al. 2000; Bethani et al. 2007; Hohenstein & Roche 2001; Kreykenbohm et al. 2002; Kweon et al. 2003; Tsui & Banfield 2000; Tsui et al. 2001; Xu et al. 2014). The ability to form stable non-cognate SNARE complexes has also been shown by biochemical studies (Fasshauer et al. 1999; Furukawa & Mima 2014; Izawa et al. 2012; Wiederhold et al. 2010; Yang et al. 1999). Promiscuous SNARE interactions could be less efficient and may not lead to productive membrane fusion, as more loose complexes would not generate enough pulling force to drive fusion. Indeed, mutational studies have shown that instabilities in the neuronal SNARE complex can lead to reduced fusion activity (Sørensen et al. 2006; Walter et al. 2010; Wiederhold et al. 2010). However, for such drastic effects, mutations would have to impair the tight fit of the core of the four-helix bundle. The question arises as to whether the exchange of helices of the same basic SNARE type can lead to comparable impairments.

In the present study, we have tested a number of non-cognate combinations for their ability to form a SNARE complex. Our biochemical analyses showed that several non-cognate combinations are tolerated, while only few combinations were unproductive. When we carried out MD simulations for these complexes, our biochemical findings were corroborated: the same non-cognate combinations were largely tolerated but some combinations led to structural problems generated by a non-optimal fit of the core residues. When we reintroduced cognate core residues, the fit improved again. Thus, most non-cognate combinations lead to no or hardly measurable structural impairments, while some significantly weaken the interaction. In other words, there is indeed a slight intrinsic selectivity, but by and large this only applies to a few non-cognate SNARE combinations. This is not surprising as the conservation pattern of the core residues does not vary greatly in the SNARE subtypes. Thus, it can not always be assumed that an exchange of a SNARE subtype impairs the tight fit of the core drastically.

Many non-cognate SNARE combinations probably do not interact in the cell, because they perform their task in different subcellular locations. The suboptimal fit of some non-cognate SNARE combinations discovered here could thus have arisen because cognate SNARE combinations have coevolved due to their physical separation in the cell. However, one must bear in mind that the sequence space for mutations of the interacting, mostly hydrophobic amino acids in the core of the SNARE bundle is not particularly large, since the assembly of the bundle must generate enough force for membrane fusion. The need to conserve their enzymatic activity thus largely prevents them from drifting apart. Note also that there is only one ATPase N-ethylmaleimide-sensitive factor (NSF) with its cofactor to dissociate the different SNARE complexes and thus activate them for the next fusion step. This also restricts the design options for SNARE complexes. Our analyses indicate that the interaction surfaces of some SNARE complex combinations have changed together, possibly only in certain eukaryotic lineages. Detecting these co-changes would require detailed covariation analyses, which take into account the evolutionary history of the interaction partners.

That SNARE proteins have limited selectivity in partner choice is exploited by some pathogenic bacteria and viruses which express SNARE proteins when they are in the host cell, potentially diverting certain transport pathways in the cell (Khalifeh et al. 2022; Neveu et al. 2020). Apparently, this can even be accomplished by proteins that have evolved SNARE-like properties, that is, they have a coiled coil region that can interact with SNARE proteins. This is referred to as molecular mimicry (Paumet et al. 2009; Shi et al. 2016).

It is interesting to note in this context that the SNARE reaction has inspired the development of synthetic fusion machines. Among other things, DNA strands have been attached to membrane anchors and inserted into the membrane of liposomes (Khvotchev & Soloviev 2022; Paez-Perez et al. 2022; Sadek et al. 2016; Seeman & Sleiman 2017). They then interact with the complementary DNA strands in another liposome to fuse the membranes. It can be measured how much energy of formation of the double helix is required for membrane fusion. Due to its chemical complementarity, DNA can be programmed like a barcode and thus generate specificity. This has great potential for liposome-based drug delivery approaches. Whether one could use the QabcR code of SNAREs studied here as a design principle as well is unclear.

To understand why there is a diverse repertoire of SNARE proteins in eukaryotic cells, we need to take a closer look at the evolutionary history of these proteins. It tells us that the various QabcR sets, which already worked in different trafficking steps between compartments in the LECA, most probably evolved from an ancestral QabcR complex that was present in the pre-LECA cell. The ancestral SNARE complex was duplicated, perhaps by genome duplications. Most probably, not only the SNARE fusion machinery but also many of the key factors interacting with the SNARE proteins such as SM, Rab, and CATCHR proteins (Jahn 2024,Koike 2022,Pelham 2001,Wendler 2001,Linial 1997) were duplicated in this way during the rise of the eukaryotic cell. It can be assumed that the existing diversity of SNARE proteins and their interaction partners reflects the increasing separation and specialization of the intracellular compartments during the early evolution of the eukaryotic cell. The resulting homologous interaction networks specialized for the various transport steps of the eukaryotic cell and the compartments gradually took on different tasks. These interaction networks contribute to specificity of SNARE complex formation. The *C*-terminal transmembrane region of the SNARE proteins probably also plays an important role in which membrane the proteins are located (Sharpe 2010). There are therefore different levels of selection and targeted activation of SNARE proteins before they engage in fusogenic SNARE complexes in the cell. These may ultimately result in only certain SNARE proteins entering a certain transport vesicle and being activated for fusion.

## Methods

### SNARE protein sequences

Sequences of SNARE proteins were retrieved from our large collection from a broad range of eukaryotes. We used about 18,000 unique SNARE protein sequences that were classified according to specific hidden Markov models (HMMs) (Kienle et al. 2009; Kloepper et al. 2007). Basically, the sequences were classified into four basic types, namely Qa-, Qb-, Qc-, and R-SNAREs. These types were in turn further subdivided into different cellular transport steps: ER (I), Golgi (II), endosomes (III), and secretion (IV). The sequence logos were generated from alignments of SNARE motifs using the WebLogo software (Crooks 2004).

### Physicochemical descriptors of side chains

Various physicochemical descriptors of amino acid side chains were selected to investigate the sequences of SNARE proteins. The descriptors included: (I) Van der Waals volume (vdW), (II) hydrophobicity (Tanford scale) (Tanford 1962), (III) flexibility , (IV) bulkiness (Zimmerman et al. 1968), (V) steric factor (also known as Graph shape index) (FAUCHÈRE 1988), and (VI) the maximal number of hydrogen bonds a side chain can establish (FAUCHÈRE 1988).

### Estimation of the available and occupied volume in the hydrophobic core of SNARE complexes

The available volume of the hydrophobic core of a SNARE complex was inferred from virtual cubicles between Cα atoms of amino acids in ’a’ and ’d’ registers of the SNARE motifs. For this, the available crystal structures of SNARE complexes were used (<u>1sfc</u>, <u>2nps</u>, <u>3b5n</u>, <u>1gl2</u>, and <u>4wy4</u>). Note that our calculation is based on the assumption that the side chains of ’a’ and ’d’ positions occupy half of the volume of each of two neighboring cubicles. To calculate the occupied volumes in the complex, we summed the average side chain vdW volume of each core layer of all four basic SNARE types, Qa, Qb, Qc, and R, across all eukaryotes. This was carried out using all collected SNARE sequences. Additionally, averaged volumes distributions per layer for complexes were also estimated for SNARE subtypes involved in different trafficking steps.

### Testing the ’QabcR rule’ of SNARE complexes

The 24 different SNARE proteins of *Saccharomyces cerevisiae* were combined to form hypothetical complexes. Our HMM classification divides the different baker’s yeast SNARE proteins into four transport steps: 4 type I, 6 type II, 8 type III, and 6 type IV. The latter type contains 2 Qbc SNAREs, resulting in a total of 8 different SNARE motifs for type IV. For each SNARE type or trafficking step, we created all possible combinations of four SNARE motifs, including combinations with multiple identical motifs and calculated the occupied volumes in each of the hypothetical complexes as described above. To see how well each of these combinations would fit into the given spindle-shaped framework of a SNARE complex (see Fig. 2B), we compared each of their layers with the volume distribution in the existing SNARE complex structures. For each layer, the likelihood was calculated with which the core of the layer would fit into the framework of the existing SNARE complexes using the general formula:

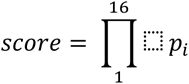

### Number of contacts in SNARE complexes

In order to find the contribution of each register in the coiled-coil structure, five available structures of different SNARE complexes (<u>1sfc</u>, <u>2nps</u>, <u>3b5n</u>, <u>1gl2</u>, <u>4wy4</u>) were selected. The interaction between amino acid residues at each register was estimated with VMD 1.92 *Hydrogen bonds* plugin. The numbers of contacts between amino acid residue pairs were plotted to establish the contribution of each register in the SNARE proteins.

### Creating and evaluating SNARE complex models using Rosetta

Four X-ray structures (<u>1sfc</u>, <u>2nps</u>, <u>3b5n</u>, <u>1gl2</u>) were selected to cover diverse subfamilies of SNARE proteins. For each of the 4 different SNARE sets, all different circular permutations of parallel arrangements in a 4-helix bundle were tested. For each bundle, 70 different permutations were generated ranging from four different helices in one complex to four helices of the same type for modeling these permutations. The SNARE motifs of the crystal structures were used as a template. For each permutated model, the sequence was threaded onto the template by using Modeller. Three independent models were generated allowing for different side chain packing. Each model was then optimized using Rosetta (https://www.rosettacommons.org/software). Each of the three starting structures was idealized and relaxed with default settings. Top three scoring structures from Rosetta relax were then optimized by using CoupledMoves protocol. Side chains and backbones were allowed to move. Top three structures from CoupledMoves were considered for further analysis. Altogether, 2520 different models were generated (4 SNARE complexes x 70 permutations x 3 initial models x 3 final models). Using Rosetta, the performance of every model was evaluated using three parameters: i) overall energy scores, ii) presence of intramolecular void volumes, and iii) atom-to-atom contacts. The parameters were then plotted for all 70 permutations for a given SNARE set.

### Protein expression constructs

All recombinant proteins were cloned into the pET28a vector, which contains an N-terminal, thrombin-cleavable His_6_-tag. The constructs for neuronal SNARE proteins from *Rattus norvegicus* have been described previously: the SNARE domain of syntaxin 1a (aa 180 –262, Syx1a), a cysteine-free variant of SNAP-25b (aa 1–206, SN25), and synaptobrevin 2 (aa 1–96, Syb2). Note that the *Rattus norvegicus* protein sequences used are identical to the ones from *Mus musculus*. Codon-optimized versions of the following SNARE domain sequences from *Mus musculus* were synthesized and subcloned into the pET28a vector (GenScript): Syntaxin 5 (aa 199-280, Syx5), syntaxin 18 (aa 229-312, Syx18), syntaxin 16 (aa 230-302 containing a single cysteine at position 231, Syx16), Sec22 (aa 128-195), Use1 (aa 174-239), and Sec20 (aa 133-198). The following Qbc-SNAREs mimicking the domain structure of SN25 were constructed: U_SNAP (Sec20, aa 133-198; Use1, aa 174-239), Me_SNAP (Membrin, aa 119-189; Bet1, aa 1-95), Go_SNAP (Gos28, 156-227; Gs15, aa 1-86), Mixed_SNAP1 (Membrin, aa 119-189; Use1, aa 174-239), Mixed_SNAP2 (Sec20, aa 133-198; Bet1, aa 1-95). Each Qbc-construct contained the linker region of cysteine-free SNAP-25 in between the Qb- and the Qc-SNARE motif. The linker has the following amino acid sequence: GLSVSPSNKLKSSDAYKKAWGNNQDGVVASQPARVVDEREQMAISGGFIRRVTND.

### Protein expression and purification

All proteins were expressed in the *Escherichia coli* strain BL21 (DE3) and purified by Ni^2+^-chromatography. After cleavage of the His_6_-tags by thrombin, the proteins were further purified by ion exchange chromatography on an Äkta system (Cytiva). The proteins were eluted with a linear gradient of NaCl in a standard buffer (20 mM Tris (pH 7.4) 1 mM EDTA) as previously described (Khalifeh et al. 2022; Neveu et al. 2020). The eluted proteins were 95 % pure, as determined by gel electrophoresis. Protein concentrations were determined by absorption at 280 nm and the Bradford assay.

### Protein interaction tests

To test possible interaction between purified recombinant proteins, protein interaction tests were used as described previously (Khalifeh et al. 2022; Neveu et al. 2020). These tests consisted of mixing different proteins together in order to find out if they formed a protein complex. In this study, equimolar amounts of the purified proteins (∼ 15 μM) were mixed and incubated overnight at 4 °C. The next day, complex formation was tested by the three-protein interaction test: SDS-polyacrylamide gel electrophoresis (PAGE), non-denaturing PAGE and size-exclusion chromatography. SDS-PAGE was carried out as described by Laemmli. Non-denaturing gels were prepared and run in an identical manner to the SDS-polyacrylamide gels, except that SDS was omitted from all buffers.

### Molecular dynamics simulations of cognate and non-cognate SNARE complexes

We generated structural models of SNARE complexes with Modeller as described above. Then we idealized and relaxed the models by using Rosetta. We selected minimum energy yielding structures for running MD simulations. For modeling, we used the same protein combinations for cognate and non-cognate SNARE complexes as used in the biochemical experiments. As cognate SNARE complexes, the ER SNARE complex consisting of Syx18 (Qa.I), Sec20 (Qb.I), Use1 (Qc.I), and Sec22 (R.I) and the neuronal SNARE complex (<u>1sfc</u>) consisting of Syx1a (Qa.IV), SNAP-25 (Qbc), and synaptobrevin 2 (R.IV) were used. To explore non-cognate complexes, we substituted the Qa SNARE of the ER complex with Syx5(Qa.II), Syx16 (Qa.III), or Syx1a (Qa.IV). Additionally, we replaced Qc SNARE of the ER complex with Bet1 (Qc.II) and also exchanged Qb and Qc SNARES of the ER complex with the Gos28 (Qb.II)/Gs15 (Qc.II) pair. In order to test the contribution of the a and d positions, we restored them for the Syx1a (Syx1a_AD) and the Gos28/Gs15 substitution (Gos28_Gs15_AD) with the original residues of the cognate ER complex. For MD simulations each system was solvated in a TIP3P water box with 12 Å of water buffer between the protein surface and the periodic box edge. Na and Cl atoms were added to neutralize the system with a concentration of 150 mM. CHARMM36 force field was used in all MD simulations with a time step of 2 fs. The simulations were performed with explicit solvent in the NpT ensemble, with a distance cut-off of 12.0 Å applied to non-bonded interactions. The temperature was maintained at 300 K using Langevin dynamics. The pressure was maintained at 1 atm using Nosé-Hoover Langevin piston. All simulations were performed with NAMD3. Three replicas were run for each system. First, each replica was minimized for 0.1 ns, gradually heated up to 300 K for 0.25 ns and equilibrated for 1 ns at 300 K. The quality of the MD simulations depends critically on the configurations visited throughout the simulation, to overcome this problem we performed 3 replicas of simulations for each system. After the equilibration, 100 ns of simulations were run for each replica. In total 300 ns of simulations were generated for each system. From 300 ns of simulations, 6000 frames were generated for analysis.

### RMSD analysis

The Root Mean Square Deviation (RMSD) was determined by using the *measure “rmsd* function” of VMD 1.9.4, by taking respective cognate complex backbone atoms, for neuronal or ER complex, as references. The backbones of the SNARE complexes with exchanges were aligned onto the respective cognate complex at each frame during the simulation and their RMSDs from the cognate complex were measured for each frame.

### CWA analysis

The helices in SNARE complexes are bent and wrapped around each other along the axis of the complex, which makes it challenging to describe their fluctuations during MD simulations. We therefore analyzed the dynamic changes of each helix pair individually – QaQb, QbQc, QcR and RQa – during simulations. To follow the fluctuations of the four helices during the simulation, we selected the Cα atoms of the ’d’ positions in the coiled coil register. They can be described as a series of squares along two adjacent helices. Due to the twisting of the helices, the four ’d’-residues do not always form a plane, but deviate from it. The deviation from the planarity was determined for each and then the cumulative deviation was calculated. Due to the fact that the complexes are in constant motion during the simulation, the CWA is given as a distribution. We noticed that during the MD simulations residues at the two ends of every chain spontaneously disengage from the α-helical structure and become more flexible. In order not to include this effect in the calculations of the CWA, the last residue in register ’d’ in every helix was excluded from the analysis. Six angles between 7 Cα atoms pairs in each helix pair were summed to produce a global wrapping parameter per helix pair, which we have designated as cumulative wrapping angle (CWA). RMSD and CWA analyses of structures were performed by a tcl script.

### Distance fluctuation analysis

To perform a comprehensive analysis and comparison of the structural traits between the cognate complex and the substitutions, we employed a distance fluctuation analysis on the Cɑ atoms of ’a’ and ’d’ registers of the four-helix bundle, which constitute the hydrophobic layers. We calculated the distances between each ’a’ and ’d’ register around each layer, generated histograms for each substitution, and then computed the Euclidean distance between the histograms of substitutions and the corresponding pair distances from the cognate ER complex. This allowed us to gain valuable insights into the variations in the hydrophobic layers’ structural characteristics between the cognate complex and its substitutions.

## Availability of data

The data that support the findings of this study are openly available in the Zenodo repository https://doi.org/10.5281/zenodo.10992340.

## Acknowledgements

This work was supported by the Swiss National Science Foundation (Grant 31003A_182732 and 310030_219549 to D.F.). We thank the Division de Calcul et Soutien à la Recherche of the UNIL for access to the university’s computer infrastructure. We thank all members of the Fasshauer Laboratory for helpful discussions.

## Author contributions

D.Y., A.H., M.D., M. D.P., and D.F. designed the study. D.Y., A.H., M.D., L.A., and D.K. performed the experiments, and analyzed the data; D.Y., A.H., and D.F. wrote the paper; and all authors reviewed and commented on the manuscript.

