## Supplemental figures for "A look beyond the QR code of SNARE proteins"

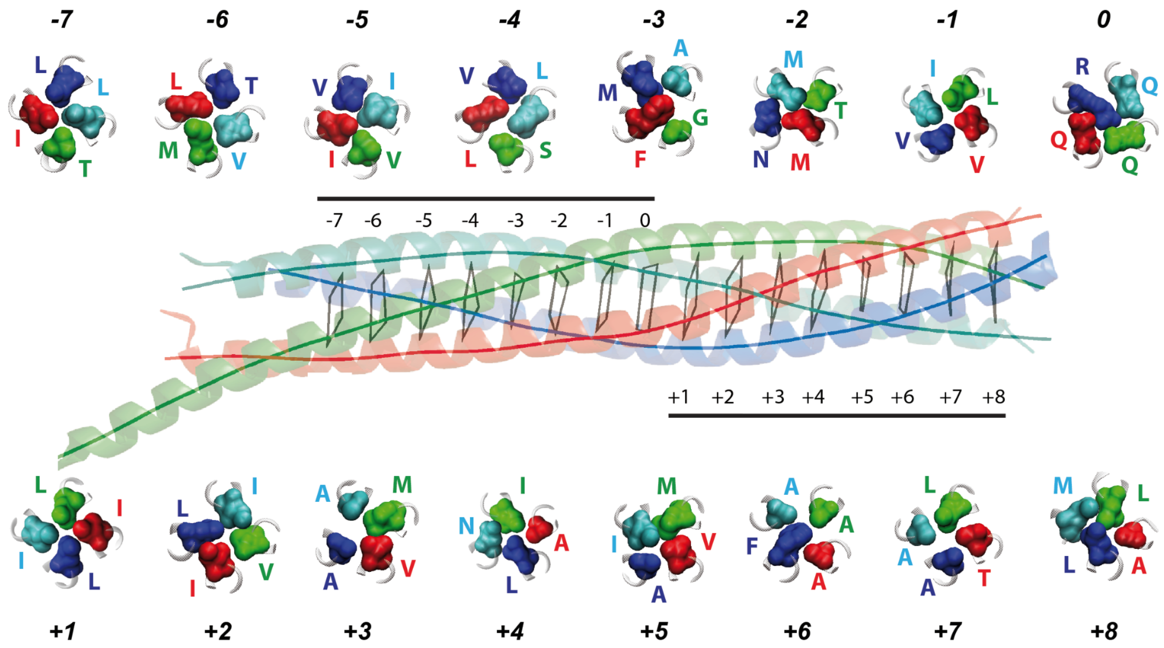


***Fig. S1. Structures of the layers in the core of the SNARE bundle.***

*As in Fig. 1, the structure of the neuronal SNARE complex (PDB: 1sfc) is shown as ribbon diagram (blue, red, and green for synaptobrevin 2, syntaxin 1a, and SNAP-25a, respectively). The 16 layers (-7 until 8) in the core of the bundle are indicated by virtual bonds between the corresponding Cα positions. The structures of the layers are shown in detail.*

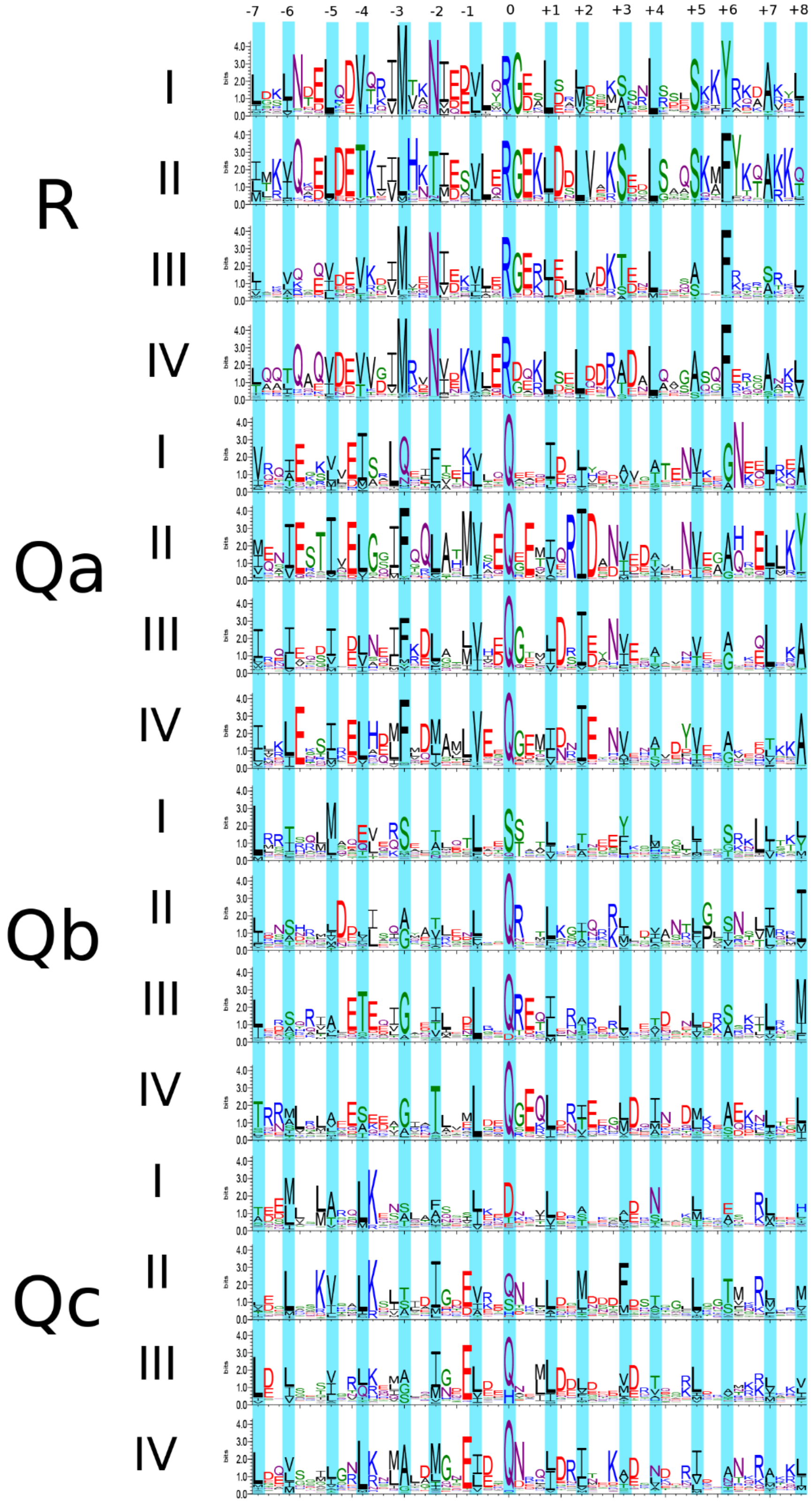


***Fig. S2. Weblogo representations of the conservation pattern of the different SNARE types.*** *The central coiled coil region of about 18,000 unique SNARE protein sequences used in our study were classified into different types based on our HMMs and aligned. From the multiple*

*sequence alignments, weblogos using the WebLogo Version 2.8.2 were generated. In detail, the height of a column in a sequence logo reflects its conservation, and the height of a letter within a column indicates its relative frequency. The residues that form the 16 coiled-coil layers are usually highly conserved. Noticeably, the weblogo representation shows that R- and Qa-SNAREs are generally more conserved than Qb- and Qc-SNAREs.*

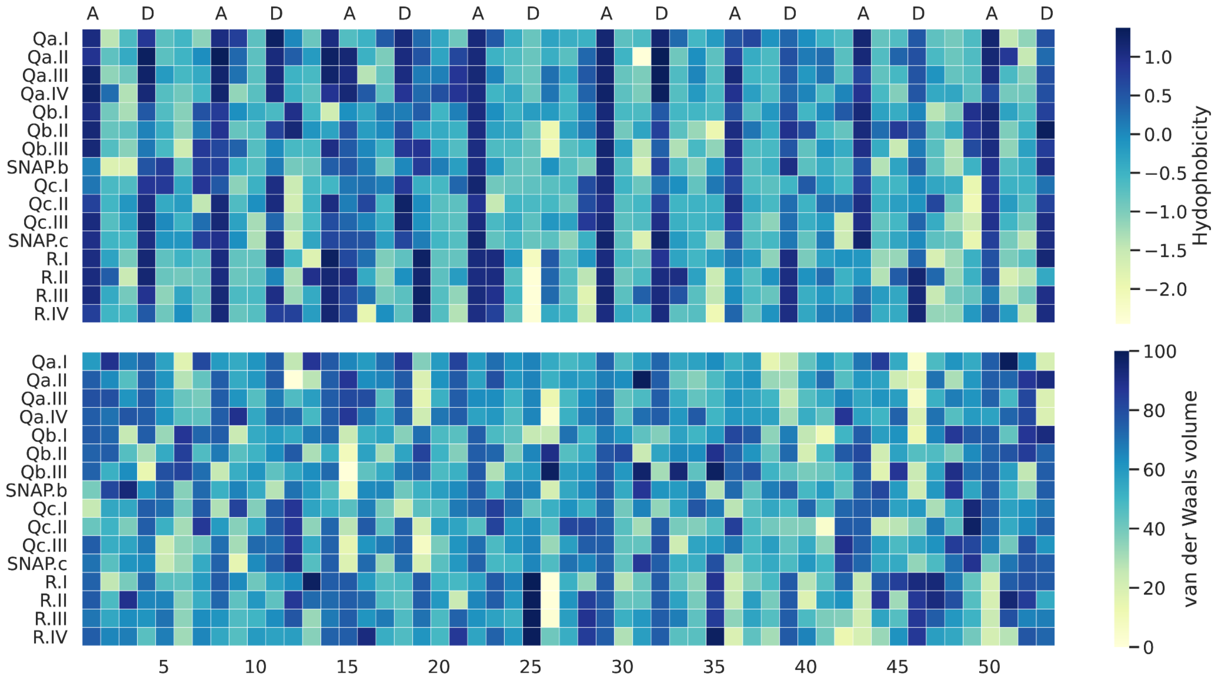


***Fig. S3. Pattern of hydrophobicity and vdW volumes in different SNARE types.***

*Multiple sequence alignments of different SNARE types (Fig. S2) were used to generate plots of hydrophobicity (upper panel) and vdW volumes (lower panel) throughout the SNARE motifs. In the hydrophobicity plots, the core layers, with the exception of the ionic 0-layer, can be recognized easily. The hydrophobicity of amino acid was according to the Tanford scale. The plot of the VdW volume of the residues in positions 'a' and 'd' is shown in Fig. 1C.*

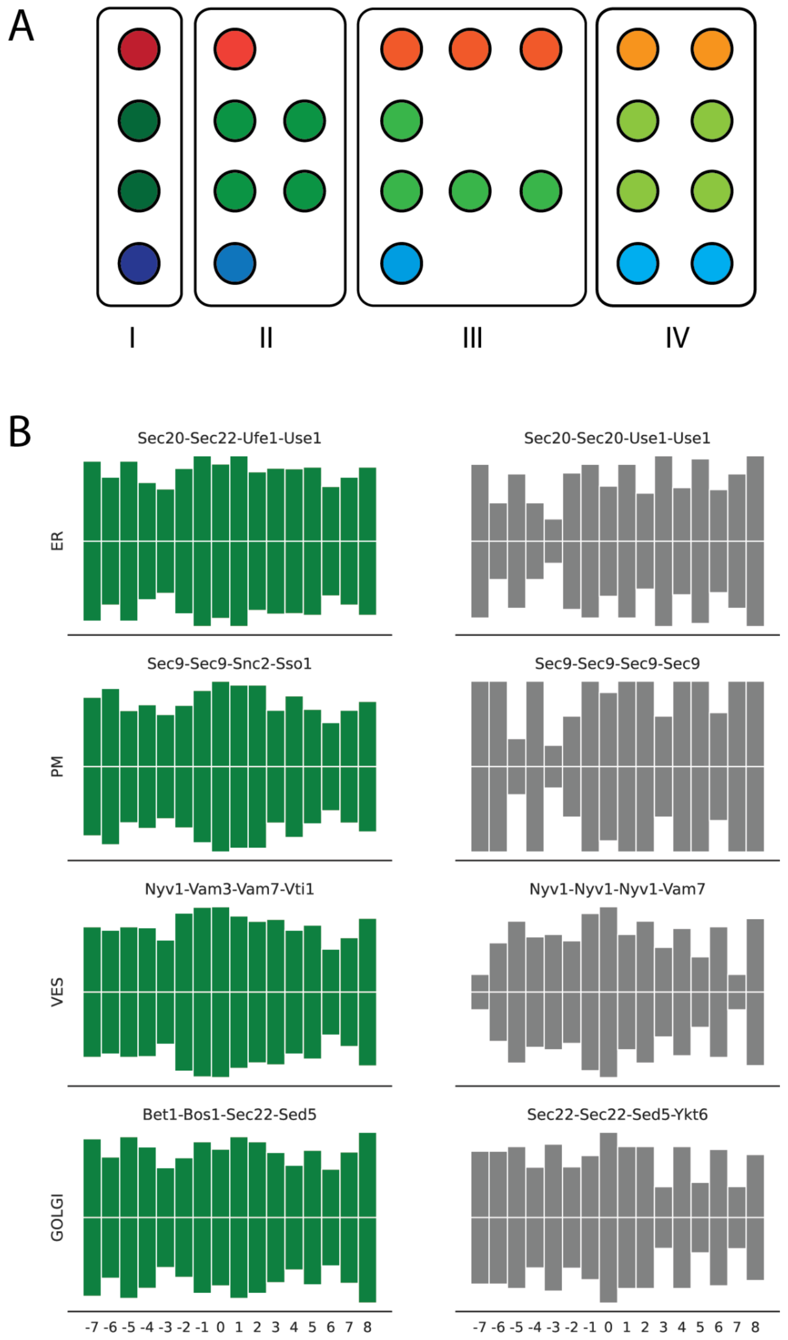


***Fig. S4. Volume distribution of yeast SNARE protein combinations.***

*The 24 different SNARE proteins from the baker's yeast were assembled in different combinations, but in the respective transport types shown schematically in (A): 4 type I, 6 type II, 8 type III, and 6 type IV. The latter type contains 2 Qbc SNAREs, resulting in a total of 8 different SNARE motifs. For each combination, the volume of the core was calculated from the VdW of the amino acids in 'a' and 'd' positions. Examples of combinations are shown schematically in (B). While the volume of established QabcR combinations shows the familiar spindle-shaped pattern, other combinations deviate significantly from this pattern.*

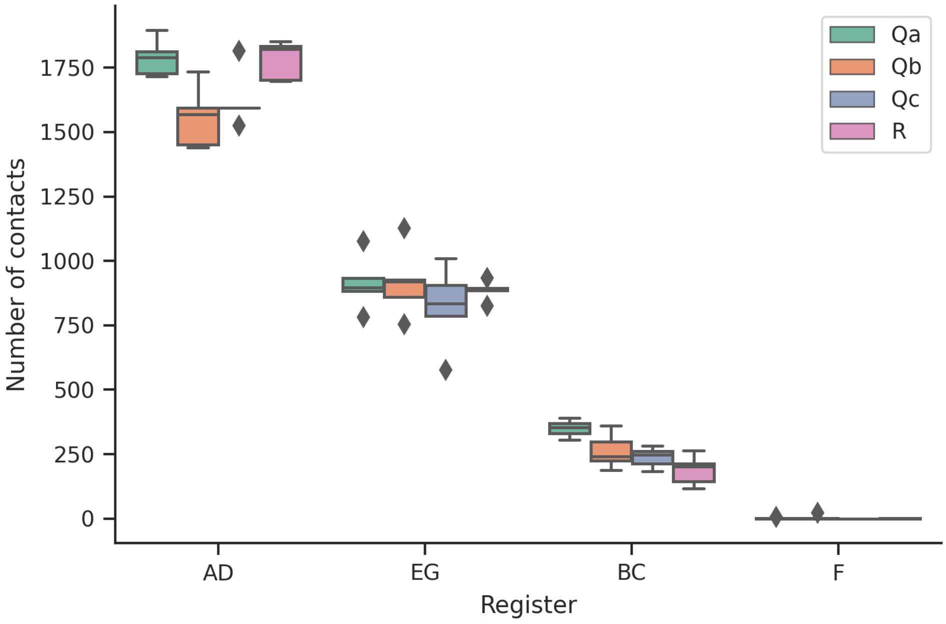


***Fig. S5. Number of contacts of residues in different positions in the heptad repeat of the SNARE bundle.***

*The number of contacts for different coiled coil registers is shown for structures of different SNARE complexes (pdb: 1sfc, 2nps, 3b5n, 1gl2).*

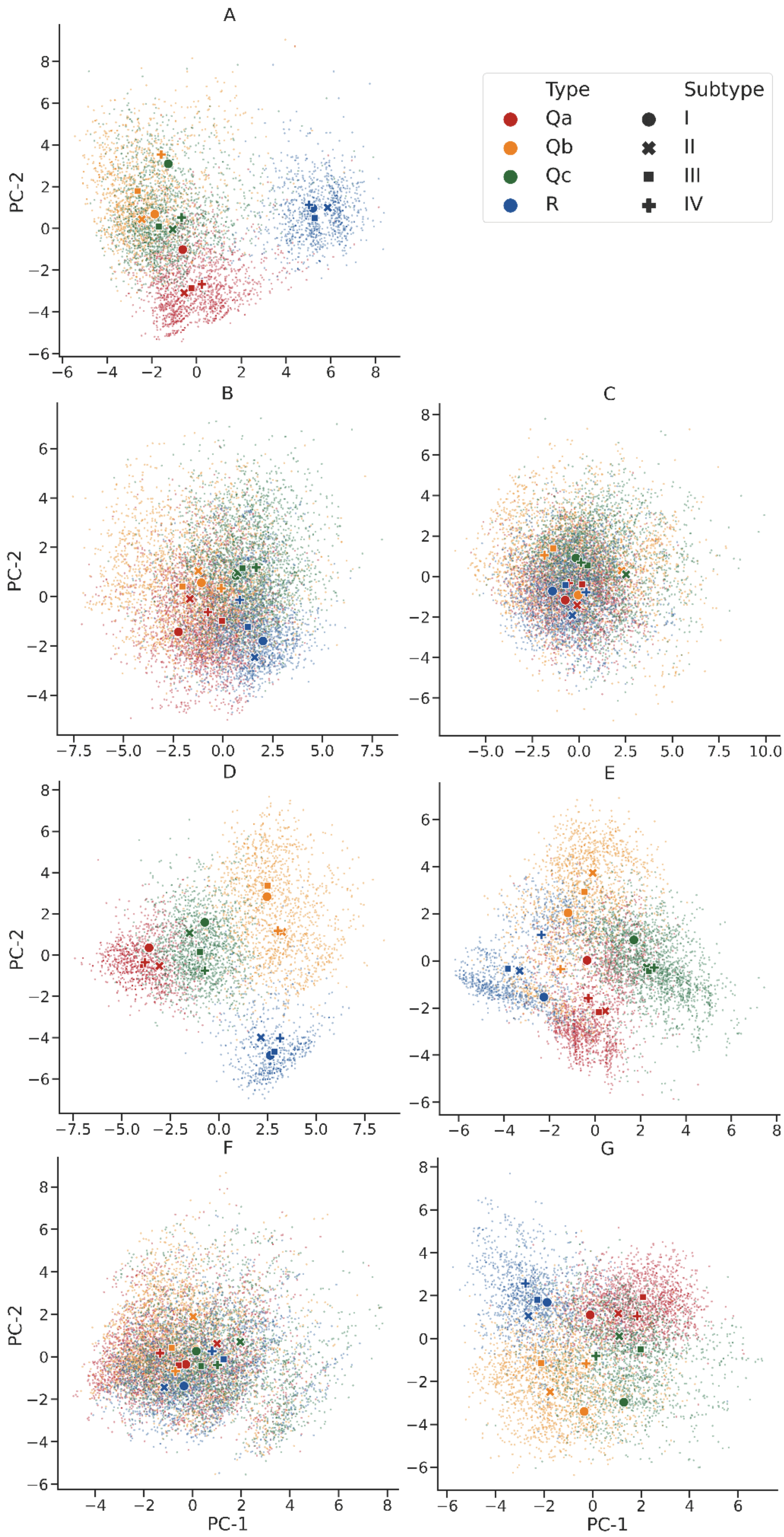


***Fig. S6. PCA of SNARE protein amino acid compositions based on the physico-chemical properties of their side chains at different positions in the heptad repeat.***

*The properties of the side chains in the core of the four-helix bundle (a and d) or between the helices (e and g) are characteristic for the respective SNARE protein type, while the side chains on the surface of the four-helix bundle (b,c, and f) hardly contribute to the distinctiveness. The calculations and plotting were carried out as described in Fig. 4.*

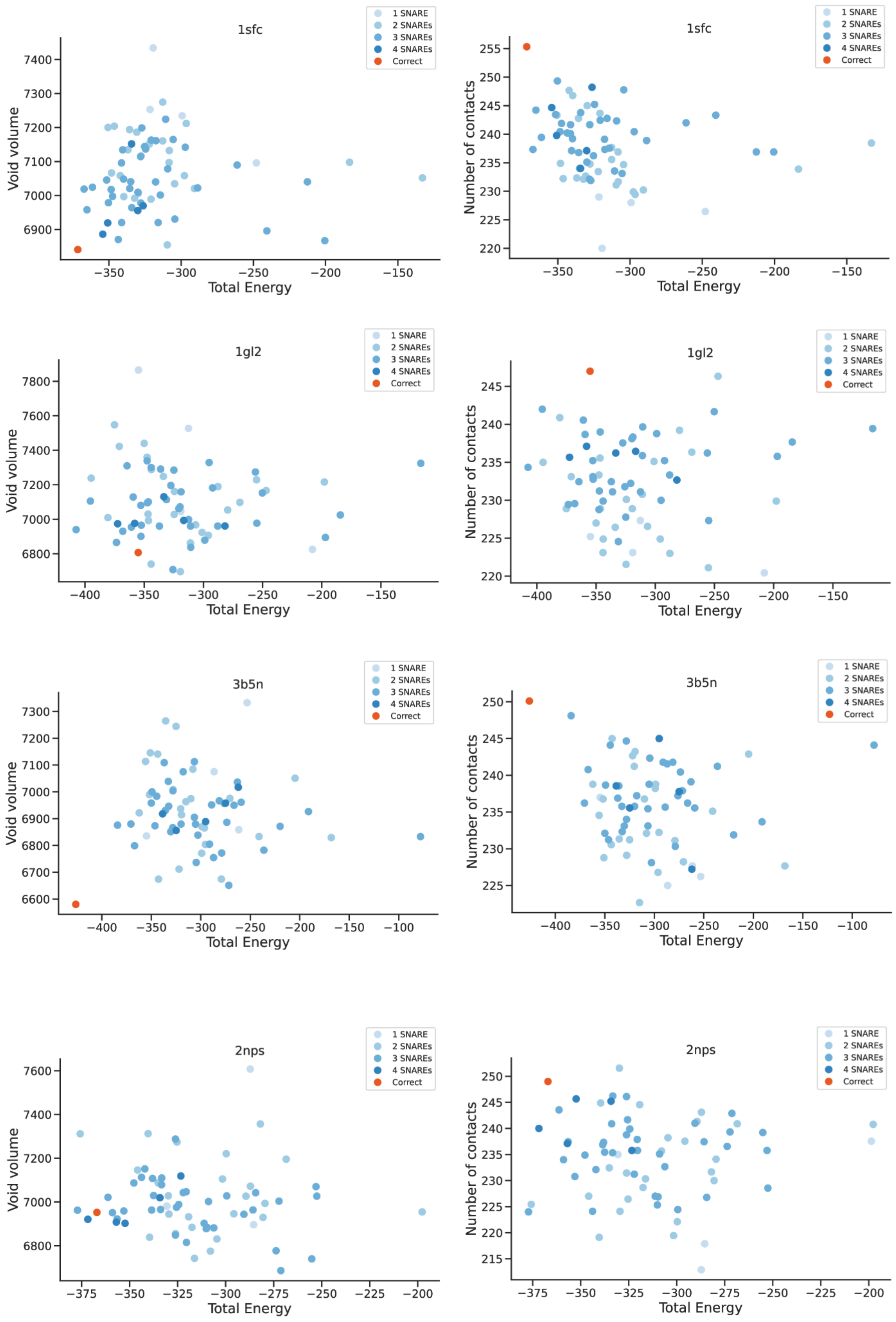


***Fig. S7. SNARE complexes in QabcR arrangement are energetically favored.***

*For different crystal structures of SNARE complexes (pdb: 1sfc, 1gl2, 3b5n, 2nps), 70 different structural permutations were modeled each and their performance was evaluated using three parameters: i) overall energy scores (in Rosetta energy units), ii) presence of intramolecular void volumes, and iii) atom-to-atom contacts. The overall energy scores were plotted against intramolecular void volumes (left) or atom-to-atom contacts (right). The referred "correct" configuration is the one which follows the QabcR rule.*

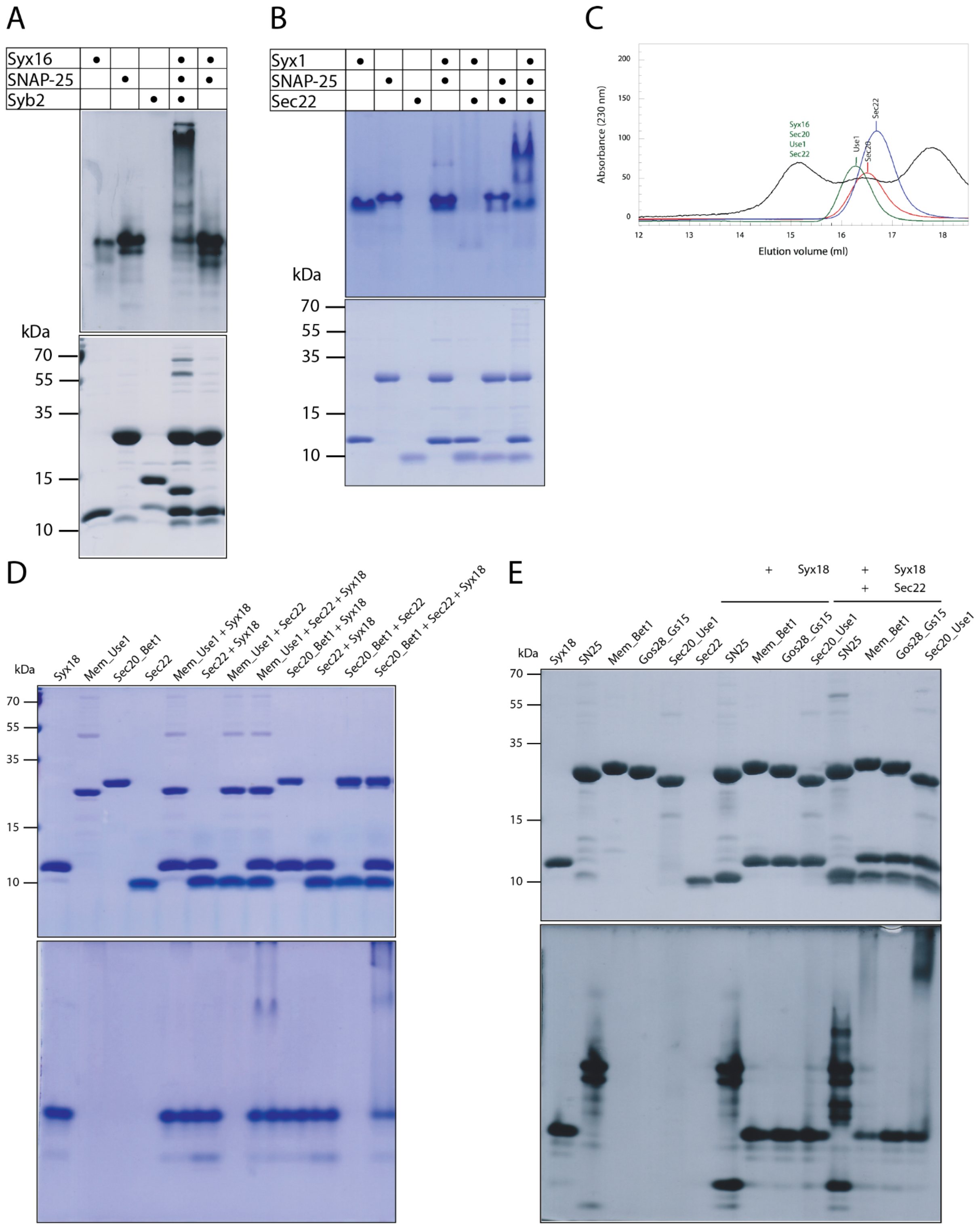


***Fig. S8. Formation of stable SNARE complexes between non-cognate SNARE proteins.***

*In all experiments, the purified monomers were mixed overnight at 4°C with equimolar ratios at ~15 µM concentration for complex formation. The mixtures and individual proteins were separated by SDS-PAGE (upper panel) and non-denaturing PAGE (lower panel)* ***(A, B, D, and E)*** *or size-exclusion chromatography* ***(C)****. When Syx1a in the neuronal SNARE complex was* *exchanged with Syx16, Syx16 was able to form a stable complex with SNAP-25 and synaptobrevin 2* ***(A)****. Note that the non-cognate complex was partially SDS-resistant. When synaptobrevin 2 in the neuronal SNARE complex was exchanged with Sec22, Sec22 was able to form a stable complex with SNAP-25 and Syx1a* ***(B)****. The ER SNAREs Sec20, Use1 and Sec22 formed a stable non-cognate complex with Syx16* ***(C)****. The ER-SNAREs Sec22 and Syx18 formed stable complexes with the chimeric Qbc-SNARE proteins Mem_Use1 and Sec20_Bet1* ***(D)*** *and Mem_Bet1 and Sec20_Use1* ***(E)*** *that contain a Qb- and a Qc-SNARE connected by the flexible linker region of SNAP-25. Note that only a weak SNARE complex band was observed when the Golgi Qbc-SNARE Mem_Bet1 was mixed with the ER-SNAREs.*

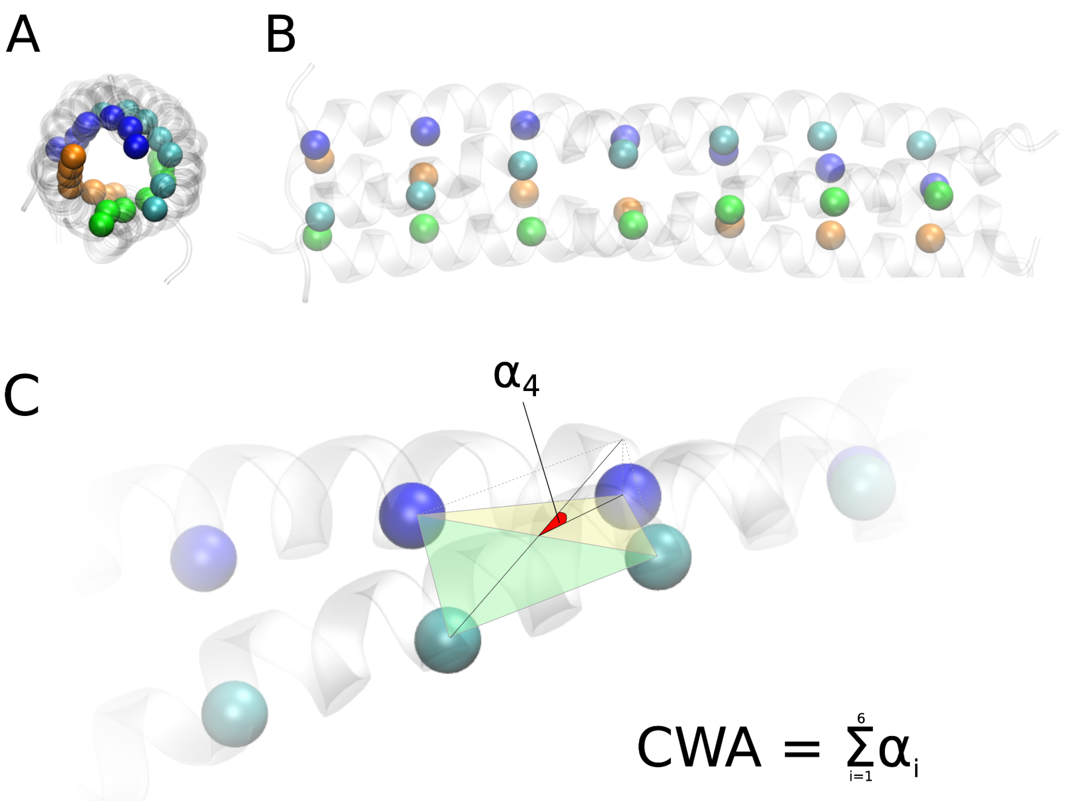


***Fig. S9. Illustration of the cumulative wrapping angle (CWA).***

*SNARE complexes are twisted four-helix bundles. Cartoon view from top* ***(A)*** *and from the side* ***(B)*** *with the 'd'-positions of the four helices shown as spheres.* ***C)*** *If two helices are parallel, the Cα of the 'd'-positions form a plane; if they wind around each other, the angle shown describes the extent to which the position of one of the Cα in space has changed.*

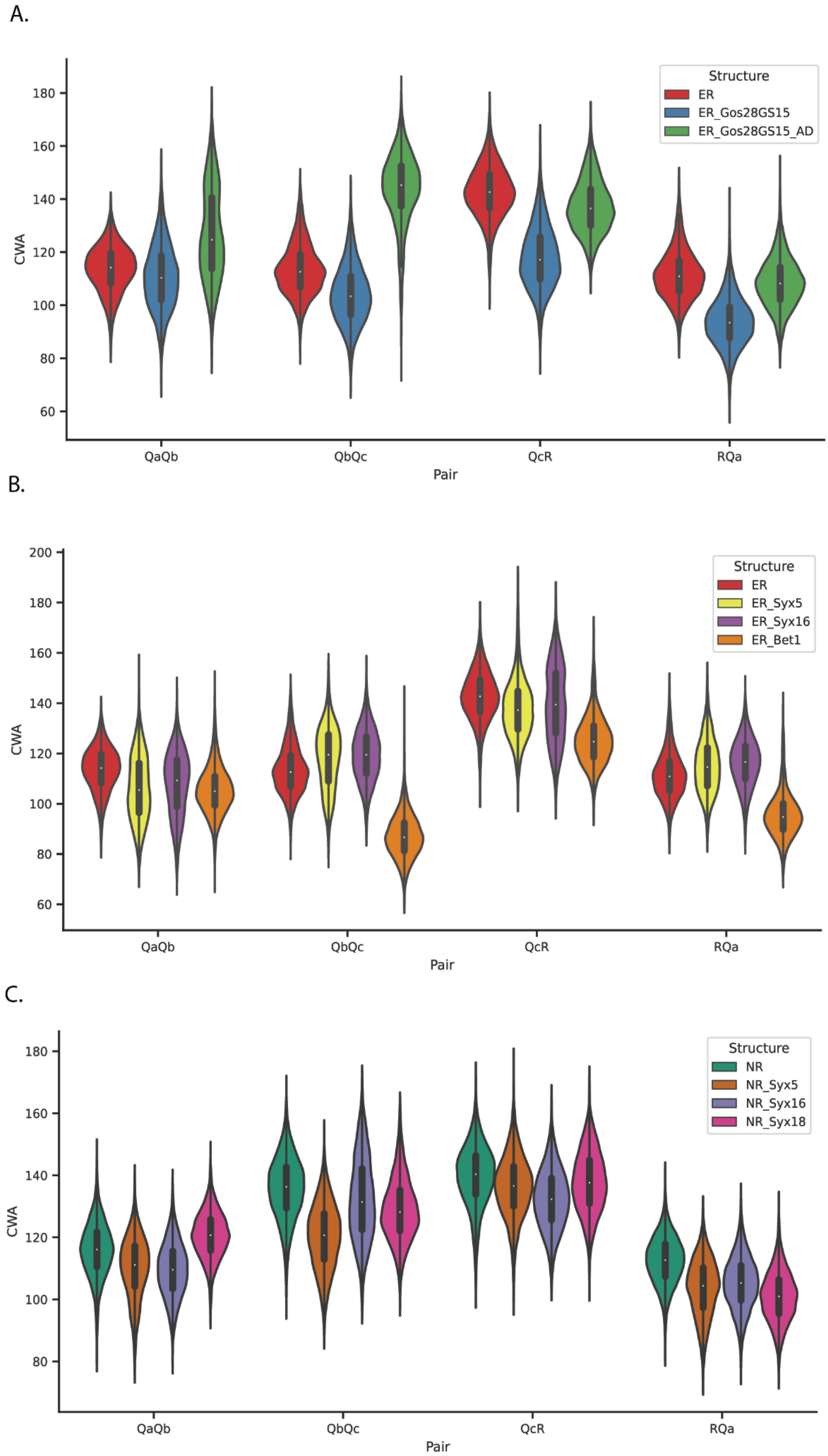


***Fig. S10. Distribution of cumulative wrapping angles for each of the four helix pairs of the ER and neuronal SNARE complexes during MD simulations.***

*In the ER complex, the cognate Qb- (Sec20) and Qc- (Use1) helices were exchanged with the Golgi SNAREs Gos28 and Gs15, respectively (ER_Gos28_Gs15). In addition, the a and d positions of Gos28 and Gs15 were swapped back with the cognate amino acids of the ER SNAREs (ER_Gos28_Gs15_AD)* ***(A)****. In another experiment, the Use1 helix was exchanged with Bet1* ***(B)****. In the ER complex, Syx18 (Qa) was exchanged with Syx16 or Syx5* ***(B)****. In the neuronal complex (NR), Syx1a (Qa) was replaced with Syx16, Syx5, or Syx18* ***(C )****.*

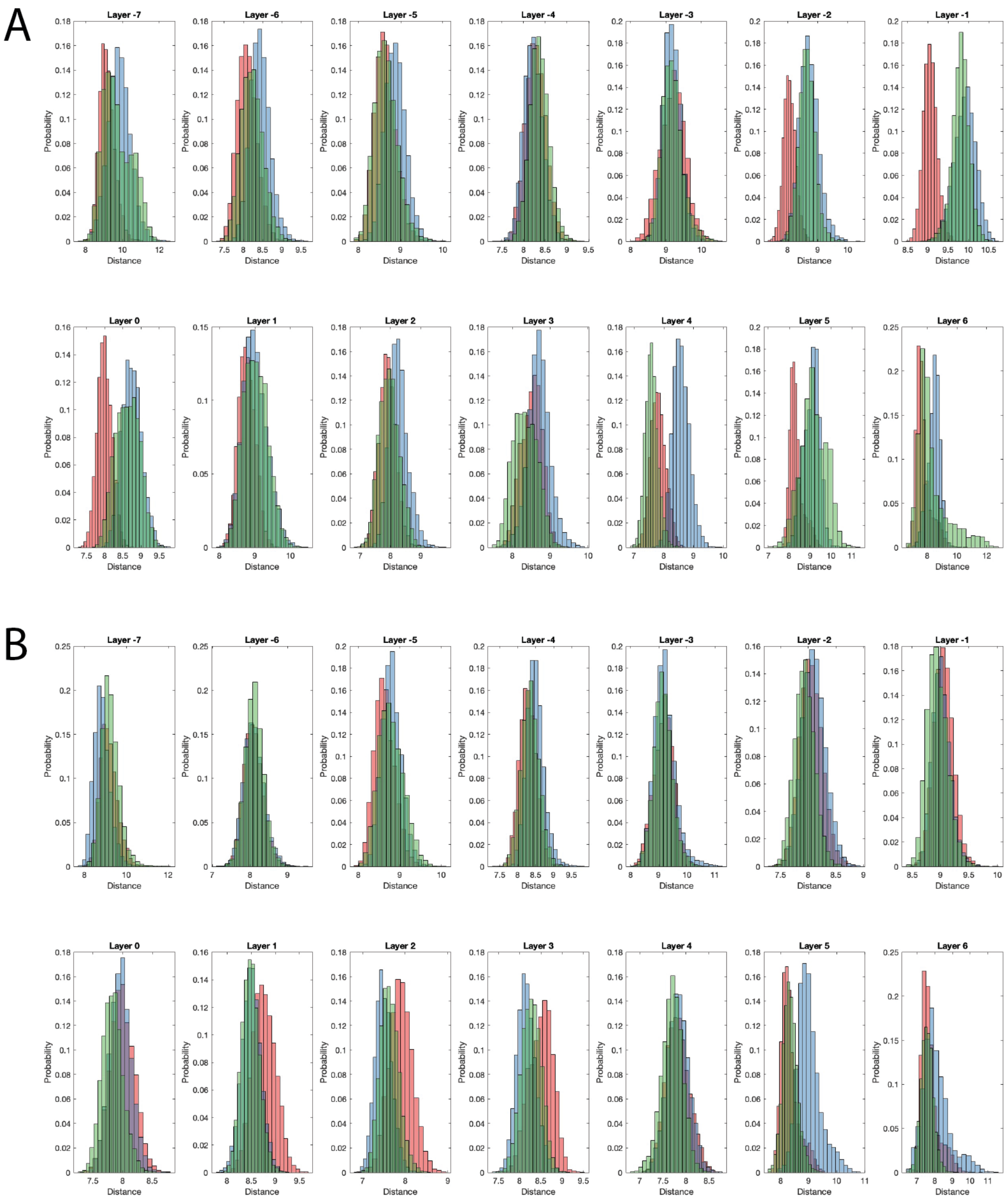


***Fig. S11. Distance distributions of layer residues during MD simulations of cognate and non-cognate SNARE complexes.***

*In the ER complex, exchanges of cognate with non-cognate SNAREs led to subtle structural problems (see also Fig. S10). A) Exchange of the Qa-helix Syx18 with Syx1a (ER_Syx1a). In addition, the 'a' and 'd' positions of Syx1a in the ER complex were swapped back with the cognate amino acids of Syx18 (ER_Syx1a_AD). B) Exchange of the cognate Qb- (Sec20) and Qc- (Use1) helices with the Golgi SNAREs Gos28 and Gs15, respectively (ER_Gos28_Gs15). In addition, the 'a' and 'd' positions of Gos28 and Gs15 were swapped back with the cognate amino acids of the ER SNAREs (ER_Gos28_Gs15_AD). Shown are the distance distributions in each layer during prolonged MD simulations. In red, the distance distributions of the cognate complex is shown, in blue the complex after exchange of a cognate with a non-cognate helix, in green the distribution after swapping back the 'a' and 'd' positions.*

***Table S1: Mean RMSD values from 300 ns of production run of each system reference to their respective cognate complex.***

| SNARE complex | Mean RMSD (Å) |
|---|---|
| Neuronal complex (NR) | 1.87 |
| NR-Syx5 | 2.14 |
| NR-Syx16 | 2.21 |
| NR-Syx18 | 1.97 |
| ER | 2.42 |
| ER-Syx5 | 2.49 |
| ER-Syx16 | 2.46 |
| ER-Bet1 | 2.81 |
| ER-Syx1a | 2.76 |
| ER-Gos28_Gs15 | 2.92 |
| ER-Syx1a$_{AD}$ | 2.05 |
| ER-Gos28_Gs15$_{AD}$ | 2.79 |
